# Novel non-transposable-element regulation patterns of KZFP family reveal new drivers of its rapid evolution

**DOI:** 10.1101/2020.04.15.041848

**Authors:** Pan Shen, Qijian Zheng, Aishi Xu, Junhua Liu, Yushan Hou, Chao Gao, Chunyan Tian, Fuchu He, Dong Yang

## Abstract

One striking feature of the large KRAB-containing zinc finger protein (KZFP) family is its rapid expansion and divergence since its formation about 400 million years ago. However, the evolutionary characteristics of KRAB domains, C2H2 zinc fingers and the full protein of KZFPs have not been fully analyzed. As for the drivers of the rapid evolution, it’s partly due to their coevolution with transposable elements (TEs). But their diverse functions besides inhibiting TEs suggest other reasons exist. Here we address these two issues by the systematic analysis of the divergence time and diversification pattern of KZFPs at three aspects and the functional analysis of the potential target genes besides TEs. We found that old-zinc-finger-containing KZFPs tend to have varied and disordered KRAB domains, indicating there are two ways of the evolution of KZFPs, including the variation of KRAB domains and the diversification of zinc fingers. Among them, the divergence of zinc fingers mainly contributes to the rapid evolution of KZFPs. Thus, we mainly focused on the functional requirements of the evolution of zinc fingers. Different from the classical regulation pattern of this family, we found and experimentally confirmed that KZFPs tend to bind to non-TE regions and can positively regulate target genes. Although most young genes tend to be with a low expression level, young-zinc-finger-containing KZFPs tend to be highly expressed in early embryonic development or early mesoderm differentiation, indicating their particular evolutionarily novel functional roles in these two processes. We further validated a young KZFP, ZNF611, can bind to non-TE region of STK38 gene and positively regulates its expression in ESCs. The emergence of new sequence in STK38 promoter may drive the evolution of zinc fingers in ZNF611. Finally, we proposed a ‘two-way evolution model’ of KZFP family.

## Introduction

KRAB domain-containing zinc finger protein (KZFP) family, the largest family of transcription factors in mammalian (Nowick et al., 2011), has 363 member genes in human genome. Generally, KZFP contains a KRAB domain and a C-terminal C2H2 zinc finger array with DNA-binding potential (supplementary fig. 1). Both C2H2 zinc finger and KRI (KRAB Interior) motif, which is the ancestor of KRAB domain (Birtle & Ponting, 2006), are old motifs, appearing widely across animals, plants and fungi. However, these two kinds of motifs did not appear in the same protein during the lengthy process of evolution until their ‘marriage’ in the last common ancestor of coelacanths and tetrapods (Imbeault, Helleboid, & Trono, 2017) about 400 million years ago. After that, the KZFP family expanded and diverged quickly especially during the evolution of mammalian.

The phenomena of the rapid evolution of KZFP family include three aspects: (1) The strong specificity of KZFP family across species. Lots of species-specific KZFP genes were reported in human, chimpanzee and other species (Huntley et al., 2006; Imbeault et al., 2017; Nowick et al., 2011; Thomas & Schneider, 2011). (2) The quick expansion of the number of KZFP member gene and C2H2 zinc finger motif in a KZFP. The number of KZFP genes (Emerson & Thomas, 2009; Looman, Abrink, Mark, & Hellman, 2002; Nowick et al., 2011) and the average zinc finger number per KZFP (Emerson & Thomas, 2009) in mammalian are much more than those in birds, reptiles and amphibians. (3) The rapid divergence of full protein sequence or C2H2 zinc finger motif among KZFP orthologs. Some studies revealed that phylogenetically specific KZFP genes are under strong positive selection (Emerson & Thomas, 2009; Looman et al., 2002). The specificity of the binding sequence is depended mainly on three key amino acids within each zinc finger (at positions 6, 3 and -1 of the C2H2 helix) (supplementary fig. 1), and the amino acid at position 2 which contacts the secondary strand (Ecco, Imbeault, & Trono, 2017; Emerson & Thomas, 2009). It was revealed that the key amino acids in C2H2 zinc fingers were also of great diversity among different species (Emerson & Thomas, 2009; Hamilton et al., 2006; Imbeault et al., 2017). In a word, KZFP family expanded and diversified rapidly during the evolution of mammalian lineage.

There are numerous studies focusing on the causes about rapid evolution of KZFP family. The previous studies found that KZFP can inhibit endogenous retroelements during early embryonic development (G. Wolf et al., 2015) and embryonic stem cells (D. Wolf & Goff, 2009), even in adult tissues (Ecco et al., 2016). Furthermore, a lot of data indicated that there is a coevolution between zinc fingers in KZFP family and transposable elements (TEs) (Ecco et al., 2017; Jacobs et al., 2014; Thomas & Schneider, 2011). There are two co-evolution models: (1) the arms race model (Jacobs et al., 2014), in which KZFP proteins unremittingly suppress the invasion of rapidly mutating TEs via changing the key amino acids of zinc fingers; and (2) the domestication model (Ecco et al., 2017; Pontis et al., 2019), in which KZFPs regulate domestication of TEs instead of restraining the transposition potential of TEs. Thus, the regulation of TEs is an important driver for rapid evolution of the KZFP family. However, KZFPs also have many other functions besides repressing TEs (Baudat et al., 2010; Grey et al., 2011; Grey, Baudat, & de Massy, 2018; Guo et al., 2009; Kang et al., 2012; X. Li et al., 2008; Patel et al., 2018; Riso et al., 2016; Takahashi et al., 2019; D. Yang et al., 2013), suggesting the rapid evolution of zinc fingers in KZFP family may be also related to their special functions besides repressing TEs.

Here, we focused on the deep and systematic analysis of the evolutionary characteristics of KZFP family and the reasons for the rapid evolution of this family. By identifying the evolutionary features of the KRAB domains, zinc fingers and the full protein of KZFPs, and analyzing the functions of target genes besides TEs, this study provides new understandings about the rapid evolution of KZFP family.

## Results

### Comparison of the divergence time of the full sequence, KRAB domain and zinc fingers in KZFPs

In order to fully describe the evolutionary characteristics of KZFP family, the evolutionary divergence time of each member of the human KZFP family was calculated. For each KZFP, the full protein sequence, KRAB domain and the key amino acids of zinc fingers were searched for its orthologs across species and the divergence time was determined according to the last common ancestor of the species containing its orthologs (see Method section for detail). Then the distribution pattern of these three types of divergence time was explored. Firstly, by directly comparing the divergence time distribution of the three type of divergence times, we found that, in general, the divergence times of zinc fingers are significantly later than the other two types of divergence times (fig. 1A, supplementary fig. 2A). Subsequently, we compared the difference value of divergence time between any two types of divergence times of each KZFP. KRAB divergence time is much earlier than the divergence time of full protein sequence in 40.7% of KZFPs, and is later in 31.6% of KZFPs (fig. 1B). Zinc finger divergence time is later than the divergence time of full protein sequence in 49.0% of KZFPs and is later than KRAB domain divergence time in 59.1% of KZFPs. On contrast, the divergence time of zinc finger is earlier than the divergence time of full protein sequence only in 4.8% of KZFPs and is earlier than KRAB domain divergence time only in 10.2% of KZFPs. These results showed that about half of KZFPs are much younger when they are evaluated by the divergence time of zinc fingers than by that of the full protein sequences or KRAB domains. The more newly emerged zinc fingers reveal that the rapid evolution of KZFPs in mammalian is mainly reflected in the rapid evolution of zinc fingers of them.

**Fig. 1.**
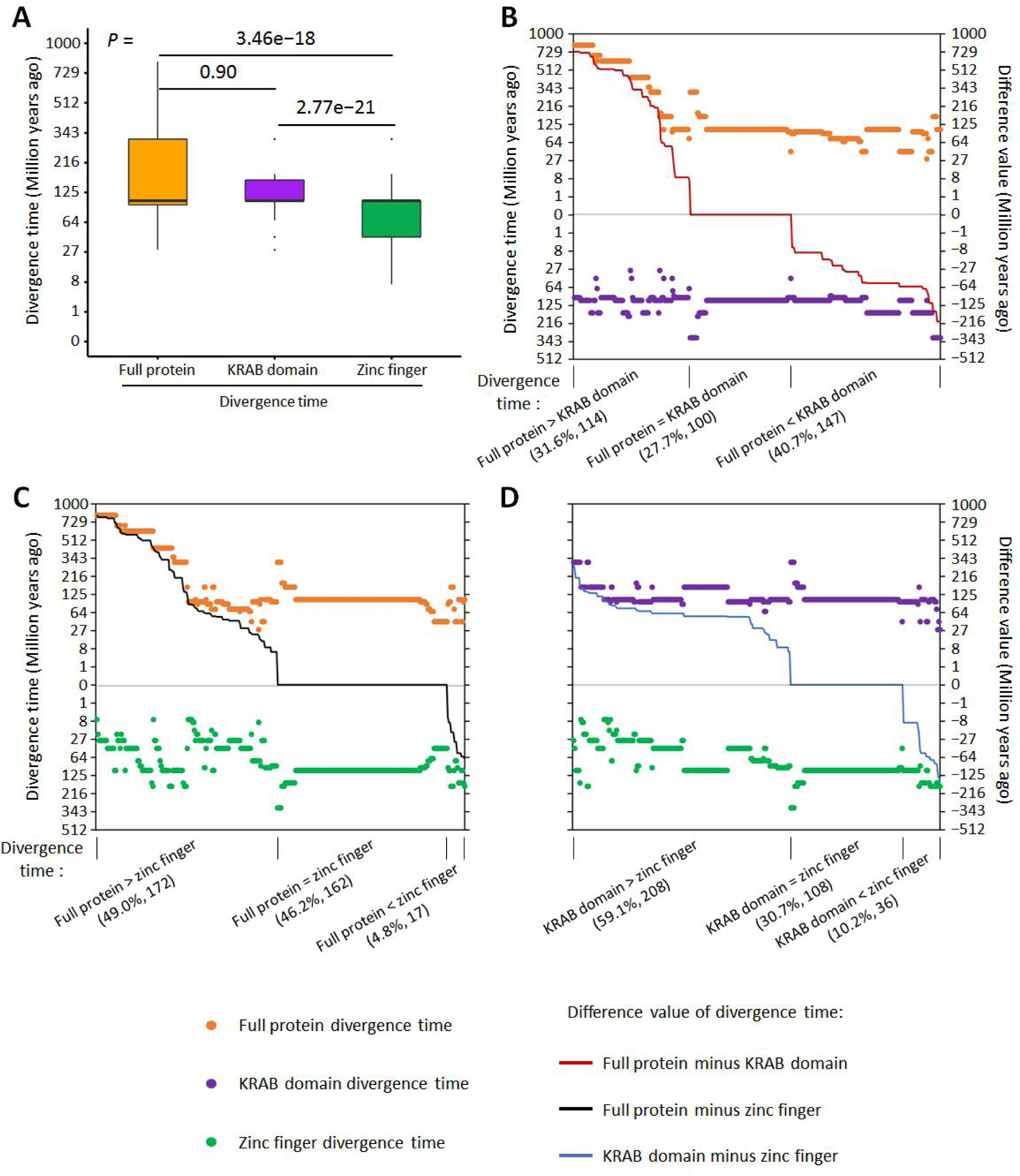
Comparison of divergence time of full protein sequence, KRAB domain and zinc finger in KZFPs. (*A*) Box plots of the divergence time of full protein sequence, KRAB domain and zinc finger in KZFPs. The values of upper and lower quartiles are indicated as upper and lower edges of the box, and the median values of median are indicated as a bar in the box. The differences of divergence time among gene, KRAB domain and zinc finger are examined by Mann–Whitney U test. The corrected P values are shown in the top of the panel. *(B-D)* Comparison of the three types of divergence time for each KZFP. In each subgraph, there are two dots for every KZFP, representing two types of divergence time. One is above the zero line and the other one is below it. The difference values of the two types of divergence time are shown as broken lines in each subgraph with the right axis as the reference scale. According to the comparison results, KZFPs are divided into three classes for each type of comparison. The numbers and percentages of KZFPs in each class are shown in brackets on the label of X-axis.

Interestingly, the three types of divergence time are all the most at the grade of Eutheria (105 million years ago, Mya). There are 133, 188, 155 KZFPs belonging to this grade of divergence time of the full protein sequence, KRAB domain and zinc finger respectively (supplementary fig. 2A&2B). Their intersection and union are 72 and 258 respectively. This result indicates that there are a large number of KZFPs originating in the common ancestor of eutherians. This may be related to the special traits in this clade. Compared with the other mammalians, such as marsupials and monotremes, eutherians have relatively long gestation periods during embryonic development (Behringer, Eakin, & Renfree, 2006), in which the KZFPs with the age of ‘Eutheria’ may play important roles (Nowick, Carneiro, & Faria, 2013).

### The diversification pattern of KRAB domains and zinc fingers in human

The rapid and frequent divergence of the sequence of KZFPs led to the diversification of KZFP members in the current existing species. To characterize the outcome of the evolution of KZFP family in human genome, the diversification patterns of the KRAB domains and the key amino acids of zinc fingers were analyzed. Through multiple sequence alignment of all human KRAB-A box sequences, we found that a cluster of 35 KZFPs have variant KRAB domains compared with other KZFPs (fig. 2A). We then supposed that the KRAB domains of these 35 KZFPs may also have special structural characteristics. The structural disorder ratio of the KRAB domains was compared between these 35 KZFPs and others. As the result, these 35 KRAB domains are significantly more disordered than the other 328 KRAB domains (fig. 2B). Disordered protein domains tend to have diverse structural conformations, and then have diverse interacting partners and functions (Csizmok, Follis, Kriwacki, & Forman-Kay, 2016; Mosca, Pache, & Aloy, 2012). Thus, we inferred that the KRAB domains in this special cluster may have diverse interacting partners besides KAP-1, the universal and canonical partner of KRAB domain. This result is consistent with a previous study (Helleboid et al., 2019) which showed that the variant KRAB box may mediate non-canonical interactions. We further investigated the distribution of the divergence time of these 35 KRAB domains. Compared with all the human KRAB domains, these 35 KRAB domains tend to be old (Amniota – Mammalia) or young (Euarchontoglires – Catarrhini), but are under-represented in middle-aged class (Theria – Boreoeutheria) (fig. 2A, supplementary fig. 2C). Even so, the middle-aged KRAB domains still account for the largest proportion in this variant cluster. This phenomenon suggests that these variant KRAB domains were formed gradually, rather than concentrated in a specific period of evolution.

**Fig. 2.**
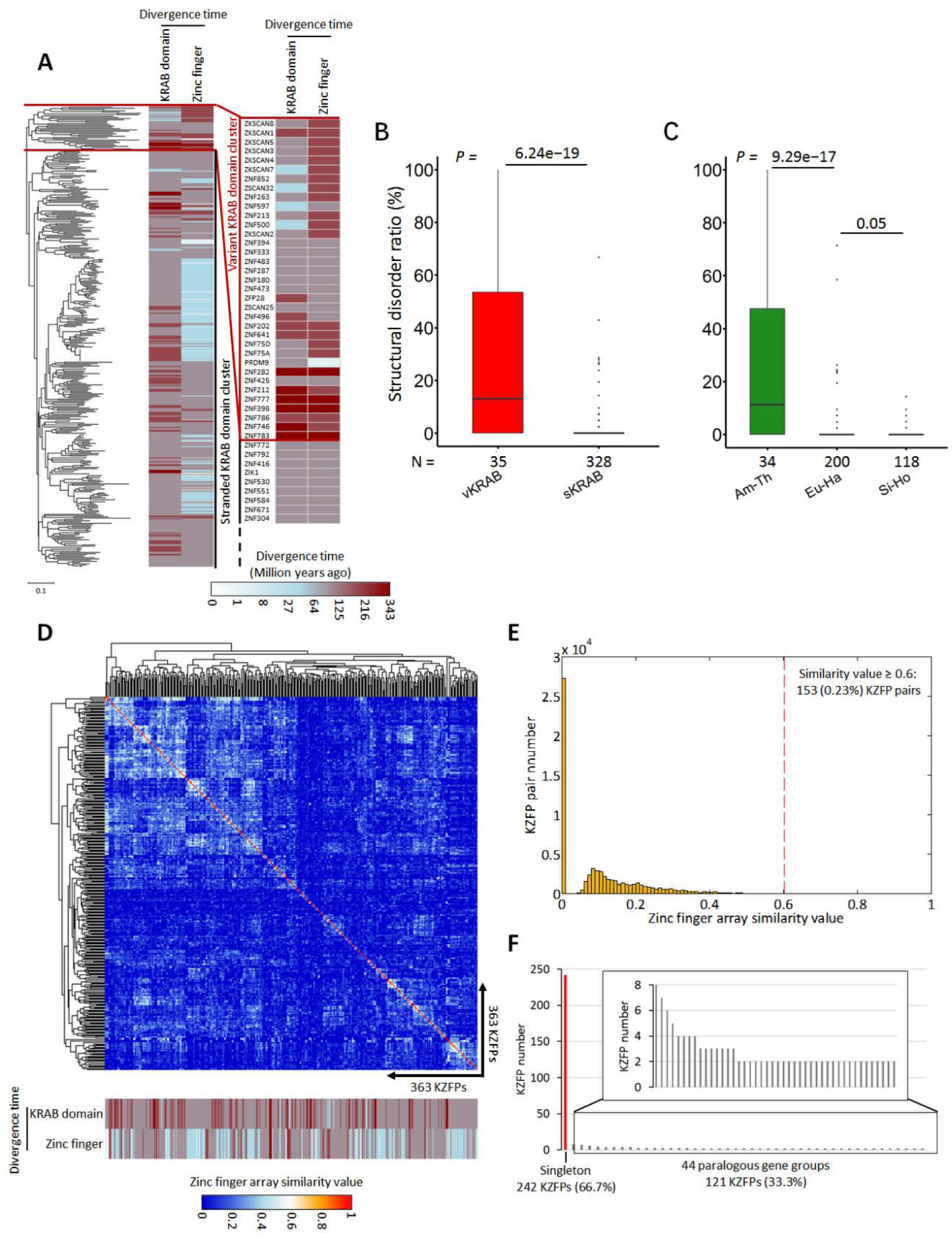
Sequence diversification of KRAB domains and zinc fingers in human KZFPs. (*A*) The phylogenetic tree of KRAB domain A-boxes in human KZFPs associated with their corresponding KRAB domain divergence time and zinc finger divergence time shown as heatmap. The variant KRAB domain cluster (right) also are displayed as a zoom-in in this figure. (*B*) The structural disorder ratio (SDR) values of the variant KRAB domains (vKRAB) and the standard KRAB domains (sKRAB). (*C*) The SDR values of KRAB domains in KZFPs with different divergence time grades of zinc finger. (*D*) The similarity values of the key amino acids in zinc fingers of KZFPs. The proportions of the similar zinc fingers (see Method for detail) in each pair KZFPs are defined as the similarity values (0-1) shown in the heatmap. As the number of zinc fingers in two KZFPs can vary significantly, we display the average similarity level. The divergence time of KRAB domain and zinc fingers of each KZFP are shown as heatmap in the bottom of the panel. (*E*) The histogram of similarity values of the key amino acids in zinc fingers of KZFPs between each KZFP pairs. The KZFP pair number and the percentage with similarity values ≥ 0.6 are shown in the figure. (*F*) The gene number and percentage of paralogous gene and singleton in KZFPs. For box-plots (*B-C*), the values of upper and lower quartiles are indicated as upper and lower edges of the box, and the median values of median are indicated as a bar in the box. The differences of SDR values between different categories are examined by Mann–Whitney U test. The corrected P values are shown in the top of each panel. The abbreviations in the figure: Am-Th, Amniota-Theria; Eu-Ha, Eutheria-Haplorrhini; Si-Ho, Simiiformes-Homo.

When the divergence time of zinc fingers was considered, among these 35 KZFPs, 23 of them (65.7%) contain old zinc fingers (the divergence time is over 159 Mya) (fig. 2A). If we calculated using the number of KZFPs containing old zinc fingers as the denominator, 67.6% (23/34) of KZFPs with old zinc fingers contain variant KRAB domains (fig. 2A). These two results revealed that there is a large intersection between the old-zinc-finger-containing KZFPs and the KZFPs with a variant KRAB domain. In other words, KZFPs with evolutionarily conserved zinc fingers tend to have a variant KRAB domain. This suggest there are two different ways for the evolution of KZFPs. One is the divergence of key amino acids in zinc fingers (that is, KZFPs containing young zinc fingers). 90% of the KRAB domains in these KZFPs are completely structured (fig. 2C), suggesting that they have a single and unchangeable function execution pattern and constantly interact with a specific co-factor (such as KAP-1). In another evolutionary path, the key amino acids in the zinc fingers are conserved (that is, KZFPs containing old zinc fingers). However, the KRAB domains of these KZFPs are variable and tend to be disordered (fig 2C), suggesting that their KRAB domains are versatile, interacting with multiple proteins.

Besides KRAB domains, C2H2 zinc finger motifs and the non-domain regions of KZFPs with variant KRAB domains (supplementary fig. 3A-3C) or old zinc fingers (supplementary fig. 3D & 3E) are more disordered than those in other KZFPs (supplementary result). But after all, KZFPs with variant KRAB domains or old zinc fingers are only a small part. For the entire KZFP family, KRAB domains tend to be completely structured although they are absolutely young domains (supplementary fig. 3F&3G; supplementary result). Furthermore, at protein level, KZFPs are also highly structured, suggesting that most of KZFPs are monotonous and unchangeable in structural conformation and functional mechanism (supplementary fig. 4; supplementary table 1; supplementary result).

In order to analyze the diversification pattern of zinc figures in human, the similarity of zinc fingers based on three key amino acids within each zinc finger in human KZFPs were calculated. As the result, the similarity between most zinc finger pairs are very low (fig. 2D, supplementary table 2), only 153 (0.23%) KZFP pairs have similarity value over 0.6 (fig. 2E). Furthermore, by identifying the homologous relationship of human KZFPs, we found that only 121 (33.3%) KZFPs belong to 44 paralogous gene groups, while 242 (66.7%) KZFPs are singletons (fig. 2F). These results indicate that the key amino acids in KZFP zinc fingers are diverse.

Overall, from the above results we can infer that the most remarkable evolutionary characteristics of KZFPs are as follows: although there are two ways of evolution for this family, that is KRAB variation or zinc finger variation, most of KZFPs belong to the ‘zinc finger variation’ evolution class. These KZFPs have relatively young and diverse zinc fingers, suggesting their targeting sequences may also evolved quickly, which acted as drivers for the quick evolution of zinc fingers.

### KZFPs tend to bind to non-TE regions in exon and promoter

In order to analyze the functional differences of KZFPs with different evolutionary features and explore the reasons for the rapid evolution of the KZFP family, the target gene information of KZFP should be investigated. To this end, we collected ChIP-seq or ChIP-exo data of 262 KZFPs (supplementary file). An important function of KZFPs is to bind and inhibit transposable elements (TEs), especially retroelements (Ecco et al., 2017; P. Yang, Wang, & Macfarlan, 2017), and there is a co-evolution model between KZFPs and TEs (Jacobs et al., 2014; Thomas & Schneider, 2011). To comprehensively analyze the genome-wide TE-binding tendency of KZFPs, we analyzed the KZFP binding sites in various regions of the genome. For a KZFP, if over half of the KZFP peaks binding to TEs in a region, the KZFP is identified as the KZFP tending to bind to TEs in this region and *vice versa*. Based on this standard, we found over half of KZFPs tend to bind to non-TE regions in genome (fig. 3A; supplementary table 3). And more interestingly, around 90% of KZFPs tend to bind to non-TE regions in exon, UTR and promoter (fig. 3A; supplementary table 3). However, according to the result of two previous studies, KZFPs tend to bind to TEs in genome (Helleboid et al., 2019; Imbeault et al., 2017). We found that main reason for this difference is the different versions of MACS program used in the ChIP-seq data processing (supplementary results; supplementary table 4). Furthermore, we confirmed that MACS2 used in this study is more accurate than MACS1.4 used in the previous studies (Helleboid et al., 2019; Imbeault et al., 2017), indicating that our conclusion that KZFP tend to bind to non-TE regions is valid (supplementary results; supplementary fig. 5).

**Fig. 3.**
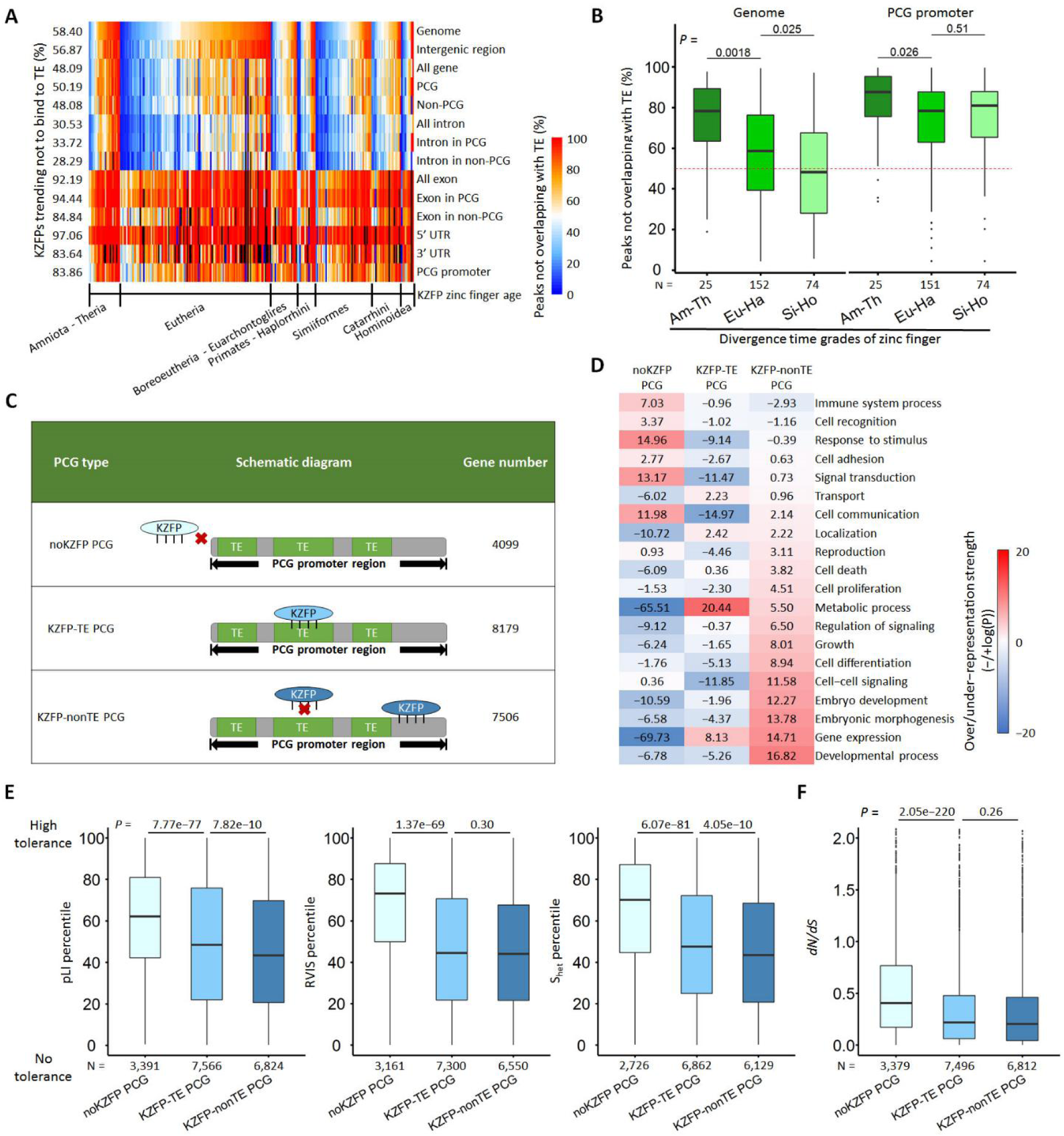
The tendency of KZFPs binding to non-TE regions and its functional significance. (*A*) Heatmap showing the percentage of the peaks not overlapping with TE for each KZFP in multiple types of genomics regions. Each row represents a type o genomic region. Each column represents a KZFP. All the KZFPs were classified into seven divergence time grades based on the zinc finger divergence time. The numbers in the left of the panel represent the percentage values of KZFPs not tending to bind to TE in each type of genomic region. (*B*) The binding bias of KZFPs with different zinc finger divergence time grades. The peaks mapping within the whole genome and PCG (protein-coding gene) promoter regions were analyzed. The red dashed line represents 50%. (*C*), The classification of PCGs into three types: noKZFP PCG, KZFP-TE PCG and KZFP-nonTE PCG. noKZFP_PCGs, the PCGs where no KZFP peak binds to their promoters; KZFP-TE_PCGs, the PCGs where at least one KZFP peak binds to TEs in their promoters; KZFP-nonTE_PCGs, the PCGs where at least one KZFP peak binds to their promoters and all KZFP peaks binding to the promoters only bind to non-TE regions. (*D*) Over- or under-representation analysis of biological processes for the three types of PCGs. The over- or under-representation strengths of each class were represented by −log (*p*) or log (*p*), respectively and were shown in the heat map. (*E*) Comparison of the tolerance to functional variation among the three types of PCGs. (*F*) the comparison of the ratio of nonsynonymous and synonymous distance (dN/dS) among the three types of PCGs. For the box plots (*B, E, F*), the values of upper and lower quartiles are indicated as upper and lower edges of the box, and the median values of median are indicated as a bar in the box.. The differences of expression width between different categories are examined by Mann–Whitney U test. The corrected P values are shown in the top of each panel.

Since there is a coevolution phenomenon between the zinc fingers of KZFP and the sequence of TE (Ecco et al., 2017; Jacobs et al., 2014; Thomas & Schneider, 2011), we assumed that there may be a tendency that KZFPs with younger zinc fingers may tend to bind to TEs. To test this speculation, we analyzed the binding bias of KZFPs with different zinc finger divergence times. Within the whole genome, it is obvious that KZFPs with younger zinc fingers tend to bind to TEs (fig. 3B & supplementary fig. 6).

We next sought to clarify the detailed functions of KZFPs which bind to non-TE regions. Based on KZFP ChIP-seq or ChIP-exo data, all of the PCGs were classified into three categories: noKZFP_PCGs, KZFP-TE_PCGs and KZFP-nonTE_PCGs (fig. 3C). We found that the biological functions of these three types of PCGs were obvious different. Interestingly, the development related processes were specifically over-represented in KZFP-nonTE_PCGs (fig. 4D), and KZFP-TE_PCGs were with higher tolerance to functional variations than KZFP-nonTE_PCGs (fig. 3E). Moreover, KZFP-TE_PCGs and KZFP-nonTE_PCGs are under higher purifying selection pressure than noKZFP_PCGs (fig. 4F). These results revealed that KZFP-nonTE_PCGs are of greatest functional essentiality among the three types of PCGs, indicating the functions regulated by KZFPs via binding to non-TE regions are very important.

**Fig. 4.**
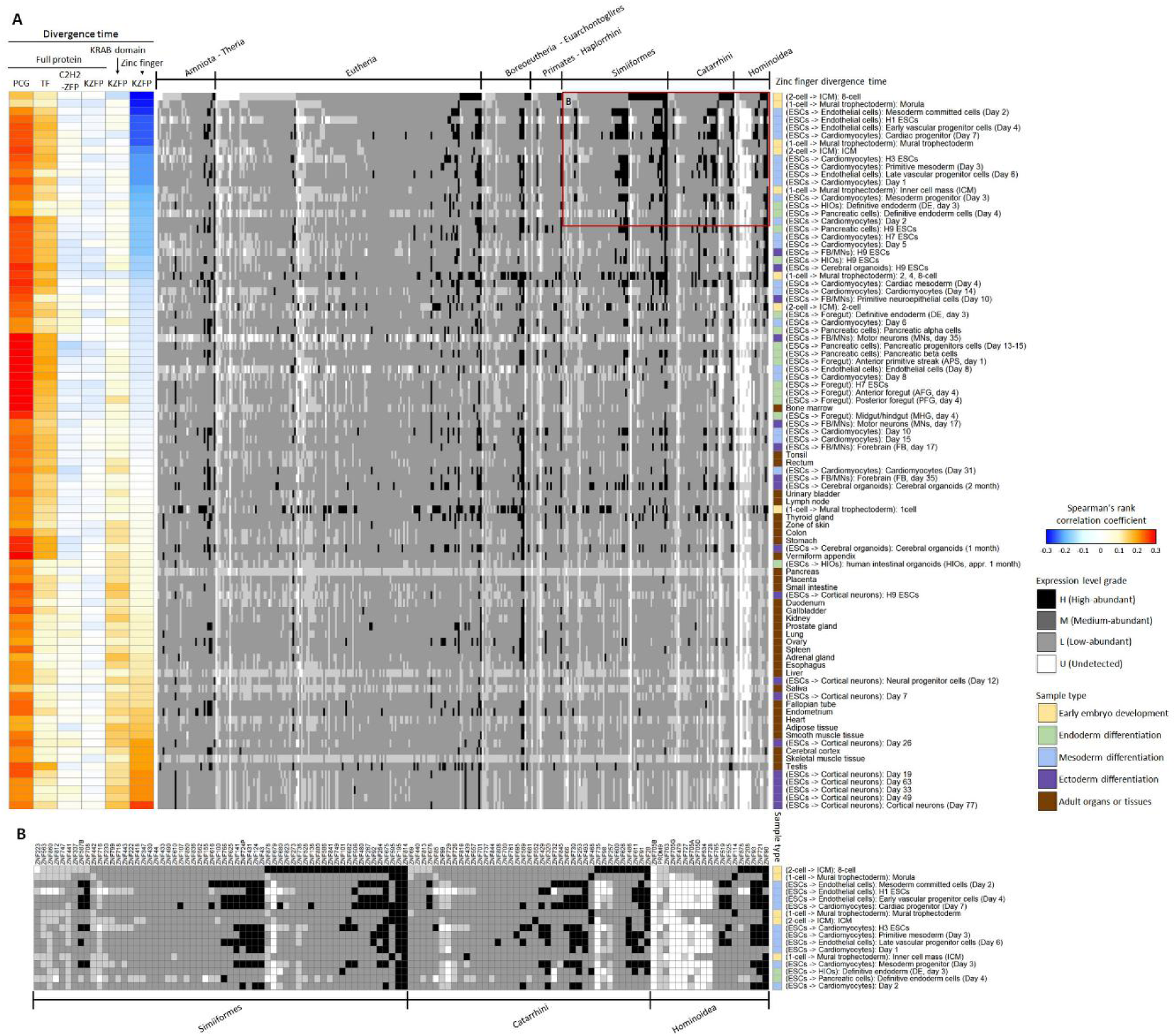
The correlation between the three types of divergence time and the expression level of KZFP genes. (*A*), left, heatmap showing spearman’s rank correlation coefficients between the divergence time of full protein sequence, KRAB domain or zinc finger in KZFPs and the gene expression level; right, heatmap showing the gene expression level which is classified into 4 grades (H, M, L, U). Each column represents a KZFP. All the KZFPs were classified into seven classes according to the divergence time of the zinc fingers. Five types of samples are marked as bars with different colors, including human early embryonic development, three directions of ESC differentiation and adult organs or tissues. (*B*) Zoom-in on the expression level of genes encoding KZFPs with young zinc fingers in early embryo development and early mesoderm differentiation.

### KZFP genes encoding young zinc fingers tend to have higher expression level in early embryonic development and the ESC differentiation into mesoderm

The preceding analyses are not related to the expression of KZFPs in certain spatio-temporal states. The investigation of the expression level of KZFPs in different spatio-temporal states is of great importance to understand their functions. In general, old genes often have higher expression level than young genes (Cardoso-Moreira et al., 2019). To verify whether the divergence time of the KZFP genes is correlated to their expression level, we explored the relationship between the evolutionary divergence time (full protein divergence time, KRAB domain divergence time and zinc finger divergence time) and gene expression level. Full protein divergence time is positively correlated with gene expression level for both TFs and other PCGs in almost all samples, and for C2H2-ZFPs in part of samples. However, this correlation for KZFPs was not significant in all samples. Interestingly, there was a negative correlation between zinc finger divergence time and expression level of KZFPs in early embryonic development and early mesoderm differentiation (fig. 4A; supplementary table 5), such as ZNF90, ZNF611 and ZNF814 (fig. 4B). In other words, in these samples, KZFPs with young zinc fingers have higher gene expression level, suggesting that younger-zinc-finger-containing KZFPs may play important roles in early embryonic development and early mesoderm differentiation.

### KZFPs can positively regulate target genes by binding to non-TE regions in endoderm or mesoderm differentiation

The data of chromatin accessibility, such as ATAC-seq data, can improve the predictability of *in vivo* transcription factor (TF) binding sites (Keilwagen, Posch, & Grau, 2019; H. Li, Quang, & Guan, 2019). Since transcription activators predominantly bound in accessible chromatin (Thurman et al., 2012), ATAC-seq data was mainly used for the prediction of TF binding sites mediating positive regulation. Interestingly, the peaks of KZFP ChIP-seq overlapping with non-TE regions were more likely to exist in open chromatin region in ESC than those overlapping with TEs (fig. 5A).

**Fig. 5.**
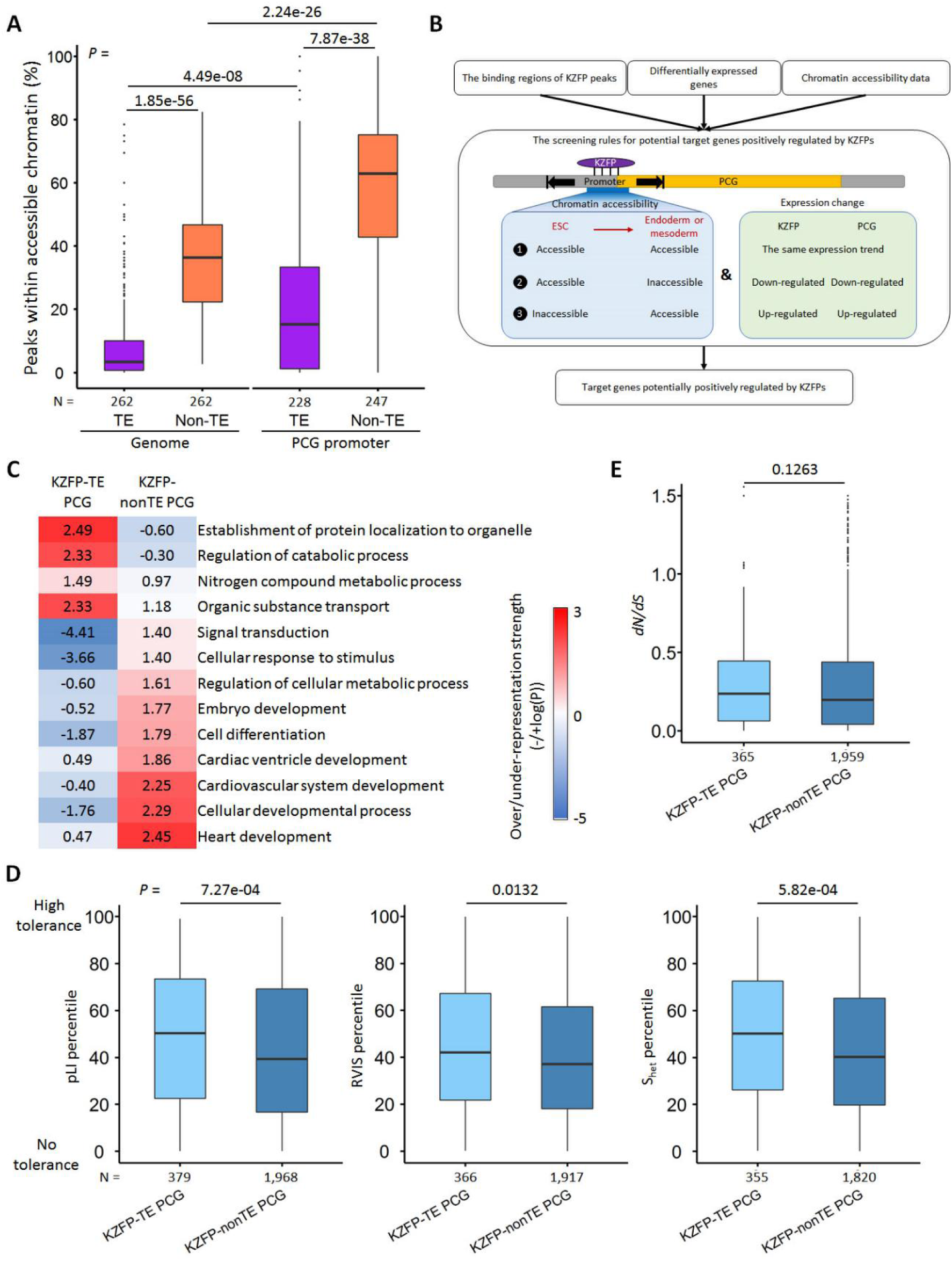
The functions of KZFP target genes in the differentiation of ESC into mesoderm. (*A*) the percentage of KZFP peaks binding to TEs or non-TE regions within accessible chromatin in the genome or PCG promoters in ESCs. (*B*) schematic diagram of the work flow for KZFP target gene screening (See method for detailed description). (*C*) the significantly over- or under-represented biological process terms for the two types of PCGs (KZFP-TE PCGs and KZFP-nonTE PCGs) in the differentiation of ESC into mesoderm. The over- or under-representation strengths of each class were represented by -log (*p*) or log (*p*), respectively and were shown in the heat map. (*D*) the tolerance to functional variants between KZFP-TEs and KZFP-nonTEs in differentiation of ESC into mesoderm. (E) the evolutionary rate of KZFP-TEs and KZFP-nonTEs in differentiation of ESC into mesoderm. For the box plots (*D&E*), the values of upper and lower quartiles are indicated as upper and lower edges of the box, and the median values of median are indicated as a bar in the box. The differences of the tolerance to functional variation and the evolutionary rate between different categories are examined by Mann–Whitney U test. The corrected P values are shown in the top of each panel.

In order to obtain high-credibility target genes of KZFPs, gene expression data (RNA-seq), ChIP-seq, ChIP-exo and ATAC-seq data were combined together to screen the target genes positively regulated by KZFPs in endoderm or mesoderm differentiation (fig. 5B). In total, we screened 4,116 target genes positively regulated by 112 KZFPs during ESCs differentiation to endoderm, and 2,490 target genes positively regulated by 76 KZFPs during ESCs differentiation to mesoderm. Of the two target gene sets mentioned above, 86.1 % and 83.8 % are KZFP-nonTE PCGs respectively. To verify the reliability of this prediction, ZNF202, ZNF383 and ZNF589 from endoderm differentiation and ZFP14, ZNF554 and ZNF565 from mesoderm differentiation were randomly chosen for the experimental validation. Interestingly, they positively regulated their target genes in ESCs but not in HEK293T cells (supplementary fig. 7A-7E; supplementary results). This could be further confirmed by the status of chromatin accessibility of the target genes in these two cell lines (supplementary fig. 7F; supplementary results). Further analysis of binding sites of 262 KZFPs in ESCs and HEK293T cells, we also found that many regions bound by KZFPs are in accessible chromatin in ESCs, while in HEK293T cells they are in inaccessible chromatin (supplementary fig. 8). These results indicated that KZFPs can play positive regulatory roles in particular biological states.

According to the traditional understanding of KZFP functions, most of KZFPs silenced TEs and also repressed the neighboring gene expression at TEs (Jacobs et al., 2014; Oleksiewicz et al., 2017; P. Yang et al., 2017). Previously, some studies also found that part of KZFPs can positively regulate target genes. For example, Schmitges *et al.* found 31 KZFPs can interact with at least one activator and it was further confirmed that ZNF554 can positively regulate the expression of target genes (Schmitges et al., 2016). Chen *et al.* found that ZFP30 positively regulated the target genes by binding to TE-derived enhancers (Chen et al., 2019). However, it is still unclear whether KZFPs may play positive regulatory roles by binding to non-TE regions. Here, we firstly report that there are a large number of target genes positively regulated by KZFPs via binding to non-TE regions in certain spatiotemporal states. These new findings expand our understanding of the functional characteristics of the KZFP family.

To understand the detailed functions of target genes positively regulated by KZFPs, we analyzed the regulatory functions of KZFP-TE_PCGs and KZFP-nonTE_PCGs in these two differentiation processes. Interestingly, we found KZFP-nonTE_PCGs specifically tend to participate in the functions closely related to the development of corresponding embryo layer in both endoderm differentiation and mesoderm differentiation, such as pharyngeal system development (supplementary fig. 9A) and heart development (fig. 5C). These results indicate that KZFPs play important roles in the differentiation of ESCs to endoderm or mesoderm via regulating target genes by binding to non-TE regions.

From the perspective of functional essentiality, KZFP-TE_PCGs have higher tolerance than KZFP-nonTE_PCGs in both differentiation of endoderm (supplementary fig. 9B) and mesoderm (fig. 5D). Moreover, KZFP-TE_PCGs are under lower purifying selection pressure than KZFP-nonTE_PCGs in endoderm differentiation (supplementary fig. 9C), while there is no significant difference in mesoderm differentiation (fig. 5E). These results further confirmed that KZFP-nonTE_PCGs are under stronger functional constraint and selection pressure, consistently indicating that binding to the non-TE regions in promoters is an important regulatory mechanism for KZFPs and the target genes that regulated by KZFPs binding to non-TE regions are more essential.

### The emergence of new sequence in STK38 promoter may drive the evolution of zinc fingers in ZNF611

As described in the preceding section, some KZFPs containing young zinc fingers highly expressed in early mesoderm differentiation (fig. 4). To validate the functional roles of these young-zinc-finger-containing KZFPs in mesoderm differentiation, the key KZFPs that may play important roles were selected and the validation experiments were conducted. Combined zinc finger divergence time, gene expression in mesoderm differentiation and the known functional annotations of KZFP target genes, ZNF611 and its target genes bone morphogenetic protein receptor, type II (BMPR2) and serine/threonine kinase 38 (STK38) were selected to do the validation experiments. BMPR2 is a transmembrane serine/threonine kinase that binds BMP2 and BMP7 in association with multiple type I receptors, including BMPR-IA/Brk1, BMPR-IB, and ActR-I (Liu, Ventura, Doody, & Massague, 1995). STK38 is a member of the protein kinase A (PKA)/PKG/PKC-like family, and STK38 can negative regulate of MEKK1/2 signaling (Enomoto et al., 2008). Both BMPR2 and STK38 are important in mesoderm differentiation, so we assumed that ZNF611 is involved in mesoderm differentiation by positively regulating BMPR2 and STK38. To test the hypothesis, we first examined the effect of ZNF611’s expression level on the mRNA level of BMPR2 and STK38 using quantitative PCR. The overexpression of ZNF611 in the ESC cells increased the mRNA level of *STK38* (fig. 6A). Similarly, when ZNF611 was knockdown with two different siRNA sequences, the mRNA level of *STK38* was decreased as well (fig. 6B).

**Fig. 6.**
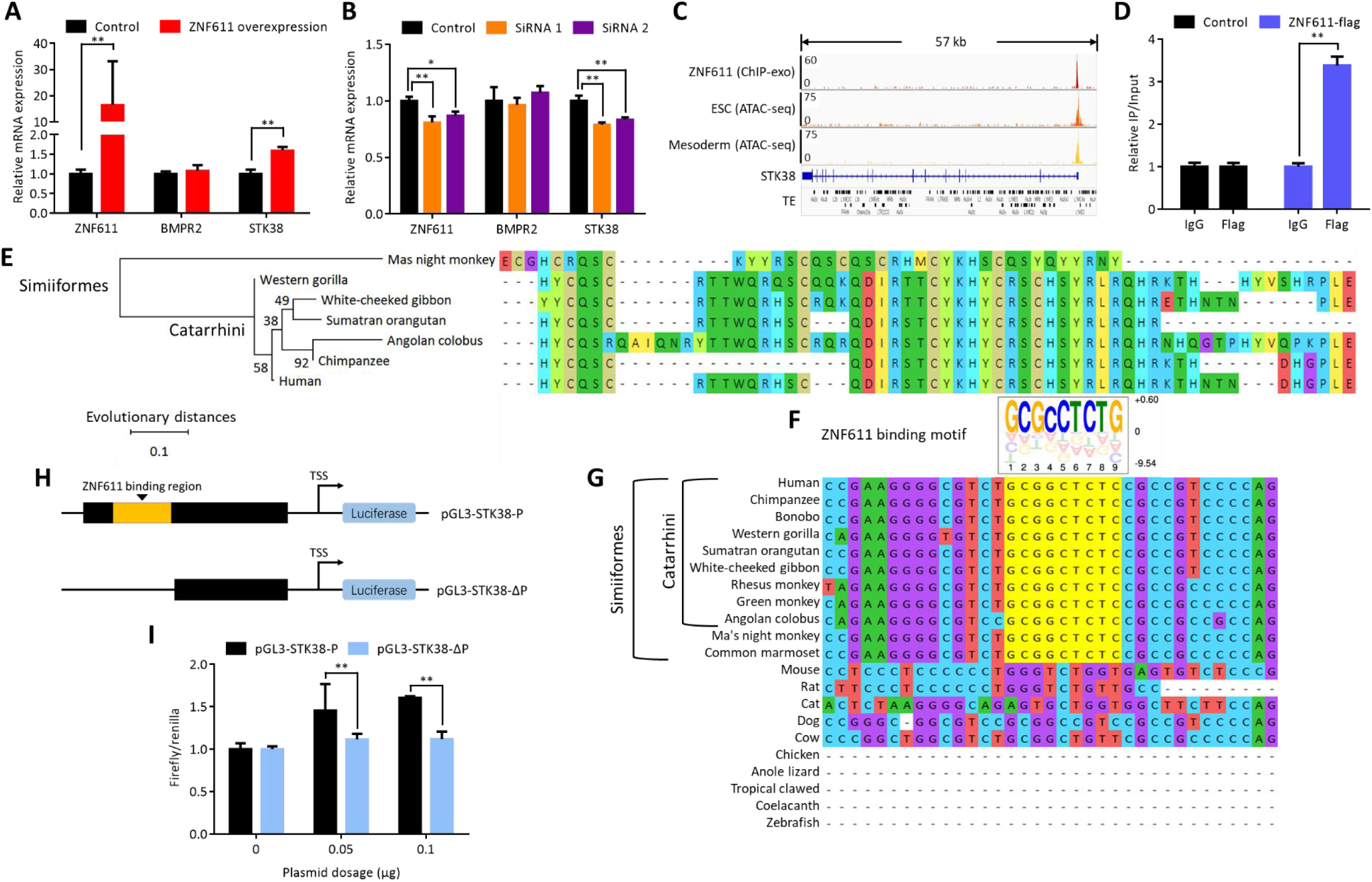
The regulatory and co-evolution relationship between ZNF611, a KZFP containing young zinc fingers, and STK38, one of its target genes in ESCs. (A) Effect of ZNF611 overexpression on the mRNA level of its target genes. Total RNA from ZNF611 transfected ESC cells was subjected to real-time quantitative PCR (qPCR) analysis. Control: empty vector. (B) Effect of ZNF611 knockdown on the mRNA level of its target genes. For panel A and B, relative mRNA levels of predicted target genes were normalized to GAPDH. The ratio values (relative expression level / the averdivergence time control value) were shown. Data are means ± SD (n = 3). The differences of expression level between control and KZFP overexpression are assessed by t-test. **: p < 0.01, *: p < 0.05. (C) Screenshot of ZNF611 binding sites and chromatin accessibility in ESC and mesoderm at genomic loci corresponding to STK38. (D) ESCs were transfected with empty vectors or Flag-tagged ZNF611 expression plasmids. After 36 h, cells were harvested and ChIP assay was performed using antibodies against IgG or Flag, and quantitative PCR was performed with primer sets against STK38 target promoters, indicating ZNF611 occupancy. Data are represented as means ± S.D. (n = 3). All data present results from three independent experiments. (E) the molecular phylogenetic tree of key amino acids in zinc fingers of ZNF611 in Simiiformes. The percentage of replicate trees in which the associated taxa clustered together in the bootstrap test (2000 replicates) are shown next to the branches. The evolutionary distances were computed using the Poisson correction method and are in the units of the number of amino acid substitutions per site. (F) ZNF611 binding motif predicted by RACDE. (G) Alignment of the STK38 promoter locus from 21 representative species. (H) Schematic representation of STK38 promoter reporter plasmids with or without ZNF611 binding site. (I) HEK293T cells were transfected with STK38 promoter reporter plasmids, and luciferase activities were measured and normalized. Representative results of three independent reporter assay experiments are shown. The data are represented as the mean ± S.D. (n = 3). All data present results from three independent experiments. pGL3-STK38-P: full construct; pGL3-STK38-ΔP: ZNF611 binding motif deletion.

Furthermore, we want to predict and verify the binding site of ZNF611 in the promoter of *STK38* gene. According to ChIP-seq data, ZNF611 can bind to non-TE region in 5’ UTR of STK38 (fig. 6C). ChIP-qPCR analysis was conducted to examining the binding of ZNF611 in non-TE region in 5’ UTR of STK38. We overexpressed the flag-tagged ZNF611 and observed an enrichment of non-TE region in 5’ UTR of STK38 in ChIPs by using flag antibodies (fig. 6D). These observations indicated that ZNF611 indeed binds to non-TE region in 5’ UTR of STK38 and positively regulates STK38 in ESCs, suggesting that ZNF611 can regulate the differentiation of ESC into mesoderm by positively regulating STK38.

We next explored whether the binding sequence in STK38 promoter drives the evolution of key amino acids in zinc fingers of ZNF611. To this end, we collected the key amino acids in the zinc fingers of ZNF611 and other KZFPs in the cluster produced by Imbeault *et al.* (Imbeault et al., 2017). Then the sequences of key amino acids were aligned and the phylogeny tree was generated (fig. 6E). We can see that the key amino acids of ZNF611 are similar with other homologous KZFPs within Catarrhini instead of other Simiiformes, consistent with the zinc finger divergence time determined by Imbeault *et al.* (Imbeault et al., 2017). We predicted the ZNF611 binding motif by RACDE (Najafabadi, Albu, & Hughes, 2015) and found that there was a binding sequence of ZNF611 in 5’ UTR of STK38 (fig. 6F). Based on the alignment of the STK38 promoter locus from 21 representative species, we found the ZNF611 binding site in STK38 are highly conserved in Simiiformes (fig. 6G). To further verify the accuracy of the ZNF611 binding site, we cloned a ∼1.2-kb DNA fragment upstream of the STK38 transcription start site (TSS) into a pGL3-basic luciferase vector (termed pGL3-STK38-P), containing the described ZNF611 binding motif. We also generated deletion lacking the ZNF611 binding motif (pGL3-STK38-ΔP) (fig. 6H). The ZNF611 overexpression dose-dependently increased the luciferase activity of pGL3-STK38-P, but not the pGL3-STK38-ΔP reporter (fig. 6I). These results suggested there was a co-evolution between the binding sequence in the 5’UTR of STK38 and the zinc fingers of ZNF611: The new sequence in STK38 appeared in the common ancestor of simiiformes, and then (about 14 million years later) the zinc fingers of ZNF611 evolved to adapt to the emergence of the new sequence. Therefore, there was a change in key amino acids in zinc fingers of ZNF611 in the common ancestor of catarrhini.

## Discussion

In this study, we tried to comprehensively characterize the evolutionary features of KZFPs by the analysis of the divergence time and diversification pattern of KRAB domains, zinc fingers and the full protein of KZFPs. We then explored the functional features of the target genes of KZFPs with different ages, so as to answer the question why KZFP family evolved so fast. We found that: (1) the rapid evolution of KZFPs in mammalian is mainly reflected in the rapid evolution of zinc fingers of them. (2) KZFPs with old zinc fingers contain variable and disordered KRAB domains. (3) Different with the previous conclusions (Helleboid et al., 2019; Imbeault et al., 2017), KZFPs tend to bind to non-TE regions, instead of TEs, particularly in exons, and promoters. (4) Different from the classical repression function of KZFPs, we found that, in certain processes, such as ESC differentiation, lots of KZFPs can positively regulate target genes via binding to non-TE regions. (5) Some young zinc-finger-containing KZFPs (e.g. ZNF611) are highly expressed in early embryonic development and early mesoderm differentiation. After experimental verification, we found ZNF611, which contains young zinc fingers, binds to non-TE region in 5’ UTR of STK38 and positively regulates its expression in ESCs, and importantly, the emergence of new sequence in STK38 promoter may drive the evolution of zinc fingers in ZNF611. These results indicate that the KZFPs containing young zinc fingers can positively regulate genes and play important roles in the differentiation of ESC into mesoderm by binding to non-TE regions, suggesting that the function of binding to non-TE regions may be one of the drivers for rapid evolution of zinc fingers in KZFPs. Moreover, STK38 was up-regulated in differentiation of human ESCs into mesoderm, while Stk38 was down-regulated in differentiation of mouse ESCs into mesoderm (supplementary fig. 10), suggesting that there is a new regulation (that is, positively regulated by ZNF611) of STK38 in differentiation of human ESCs into mesoderm. Overall, these findings show that binding to non-TE regions is one of the important ways for KZFPs to greatly contribute to the formation and evolution of gene regulatory networks.

Based on our results, combining with the conclusions of existing researches (Ecco et al., 2017; Helleboid et al., 2019), we proposed a new KZFP family evolution model: ‘two-way evolution model’ (fig. 7). The evolution of the KZFP family mainly includes two directions. One is the divergence of key amino acids in zinc fingers (that is, KZFPs containing young zinc fingers), adapting to the changes of the target sites (such as TEs reported in previous studies, or non-TE regions revealed by this study). Almost all of the KRAB domains in these KZFPs are completely structured and the whole protein of these KZFPs also are relatively highly structured, suggesting that they have a single and unchangeable function execution pattern and constantly interact with a specific co-factor (such as KAP-1). In another evolutionary path, the key amino acids in the zinc fingers are conserved (that is, KZFPs containing old zinc fingers), and their target sites are also conserved. However, the KRAB domains of these KZFPs are variable and tend to be disordered, suggesting that their KRAB domains are versatile, interacting with multiple proteins. Therefore, these KZFPs can play diverse roles.

**Fig. 7.**
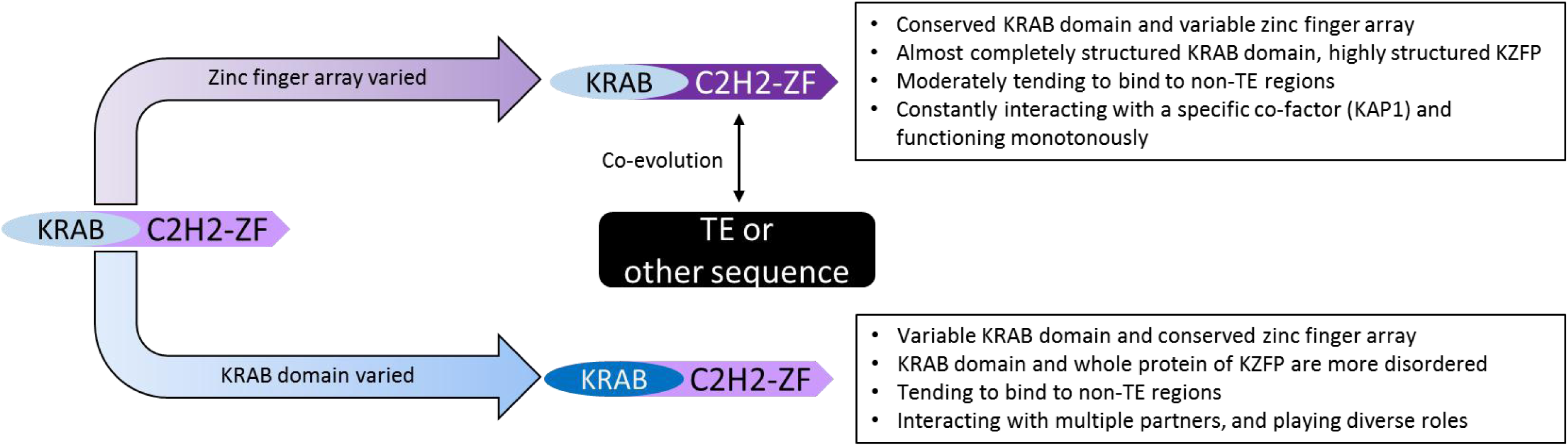
The ‘two-way model’ of KZFP family evolution. The evolution of the KZFP family is mainly divided into two directions. One is the divergence of key amino acids in the zinc fingers (zinc finger array varied). In another evolutionary path, the key amino acids in the zinc fingers are conserved, but the sequence of KRAB domain is diverged (KRAB domain varied).

## Materials and Methods

### The identification of members in KZFP family, C2H2-ZFP proteins and TFs in

#### Homo Sapiens

The *homo sapiens* protein sequences were downloaded from Ensembl release 83 (Zerbino et al., 2018), and the HMM file of KRAB domain and zf-C2H2 domain were download from Pfam v29.0 (Finn et al., 2010). The KZFPs and C2H2-ZFPs were filtered using HMMER v3.1b2 (Eddy, 2009). The validation of transcription factors was based on DBD (DNA-binding domain) transcription factor database (Kummerfeld & Teichmann, 2006) and Ref. (Ravasi et al., 2010).

### The definition of the divergence time of the full protein sequence, KRAB domain and zinc finger of KZFPs

The divergence time of the full protein sequence was inferred according to the homology information from Ensembl Compara (Herrero et al., 2016; Vilella et al., 2009). To identify the divergence time of KRAB domain in human KZFPs, protein sequences of 80 species from 80 genera in deuterostomia were downloaded from Ensembl database (supplementary table 6). KRAB domains in 80 species were identified by HMMER v3.1b2 (Eddy, 2009). BLSATP was used to calculate the homology between human KRAB domains and KRAB domains in other 79 species. The hits with the identity and the percentage of the matched sequence in query or subject sequence above 80% were selected as homologous KRAB domains. The KRAB domain divergence time of each human KZFP was determined based on the species with farthest evolutionary distance from human in all species containing the homologous KRAB domains. The divergence times of KZFP zinc finger were inferred according to the similarity between the key amino acids in zinc-fingers (Imbeault et al., 2017). Evolutionary distance between *Homo Sapiens* and other 79 species were estimated by TimeTree (Kumar, Stecher, Suleski, & Hedges, 2017). The divergence time values of full protein sequence, KRAB domain and zinc fingers of KZFPs are available in supplementary table 7.

### The phylogenetic analysis

Sequence alignments were performed using ClustalX (version 2.1) with default parameters (Larkin et al., 2007), and the phylogenetic tree (neighbor-joining tree) was constructed using MEGAX (Kumar, Stecher, Li, Knyaz, & Tamura, 2018) with default parameters.

### The zinc finger similarity between each KZFP pairs in *Homo Sapiens*

The zinc finger similarity between each KZFP pairs was calculated according to the method described previously (Imbeault et al., 2017). Briefly, similarity between two zinc finger arrays was defined as sharing the same three amino acids at the DNA-contacting residues, allowing up to one replacement by a related amino acid (as defined by Blosum62). To identify paralogs of human KZFPs based on zinc fingers: 1) for the KZFP which containing 5 or more zinc fingers, the threshold similarity score of 60% and common zinc-fingers of 5 between any two zinc-finger arrays were selected; for the KZFP which containing 4 or less zinc fingers, a threshold similarity score of 100% between any two zinc-finger arrays was selected.

### The disorder scores of proteins and domains

#### Structural disorder ration (SDR)

The longest protein encoded by each gene was selected as the representative protein for subsequent analyses. IUPred (Dosztanyi, 2018) was used to obtain the disorder score of each amino acid in a protein or domain. The disorder rate of a protein is the ratio of the number of disordered amino acids (the disorder score is greater than 0.5) to the total number of amino acids. According to SDR values, proteins can be divided into four classes: completely structured (SDR = 0), highly structured (0 < SDR ≤ 10 %), relatively disordered (10 % < SDR ≤ 30 %), and highly disordered (30 % < SDR ≤ 100 %).

#### Consecutively disordered region number (CDRN)

1) for the domains or proteins which are longer than 50 aa, a CDR consists of 20 or more consecutively disordered amino acids; 2) for the domains which are shorter than 50 aa, a region containing consecutively disordered residues exceeding 40% of all domain residues is considered to be a CDR.

### RNA-Seq data analysis

The RNA-Seq data was downloaded from GEO and ArrayExpress database (Supplementary file 1). The reads were mapped to the human genome build GRCh38 (hg38) using Salmon v0.11.0 (Patro, Duggal, Love, Irizarry, & Kingsford, 2017). Isoform expression levels were calculated as transcripts per kilobase of exon model per million mapped reads (TPMs) and read counts using Salmon v0.11.0 (Patro et al., 2017), and gene expression levels were calculated by tximport (Soneson, Love, & Robinson, 2015).

### ChIP-seq, ChIP-exo and ATAC-seq data analysis

The ChIP-seq, ChIP-exo and ATAC-seq data was downloaded from GEO DataSets and ENCODE (Consortium, 2012) (Supplementary file 1). The reads were mapped to the human genome build GRCh38 using HISAT2 (Kim, Langmead, & Salzberg, 2015). Duplicate reads, which often represent PCR amplification artifacts, were removed using SAMtools version 1.3. We used MACS version 2.1.0 (Feng, Liu, Qin, Zhang, & Liu, 2012; Zhang et al., 2008) with the ‘keep-dup all’ parameter, which is recommended for ChIP-exo as genuine reads can accumulate on a few positions following exonuclease digestion (Imbeault et al., 2017). We filtered out peaks that would meet any of these criteria: P < 1×10^−16^. The transcription start sites (TSSs) were retrieved from Ensembl release 83. The region from +2,000 bp to -500 bp of their nearest TSS were annotated as promoters (Tsankov et al., 2015). The genes whose promoters were overlapped with KZFP peaks were considered to be potential KZFP target genes.

### TE information

The TEs were obtained with the RepeatMasker track (Smit, AFA, Hubley, R and Green, P. RepeatMasker Open-3.0.1996–2010 http://www.repeatmasker.org) (Jurka, 2000) from the hg 19 and hg38 genome assembly.

### Screening process of KZFP target genes

If the promoter region of a PCG is bound by a KZFP, the PCG is considered as a possible target gene of the KZFP. Genes which were expressed at least in one stage (ESC, endoderm or mesoderm) were included in the following analysis. Differentially expressed genes from ESCs to endoderm or mesoderm were screened according to the fold changes in four data sets (up-regulated trend: The fold change is greater than 1.1 at least in three data sets; down-regulated trend: The fold change is less than 0.91 at least in three data sets). Chromatin accessibility data in ESCs, endoderm and mesoderm were used to filtrate the KZFP target genes: 1) if a KZFP peak bind to a promoter of PCG with accessible chromatin in both ESC and endoderm or mesoderm, the expression trend between KZFP gene and PCG should be same; 2) if a KZFP peak bind to a promoter of PCG with accessible chromatin only in ESC, both KZFP gene and PCG should be down-regulated; 3) if a KZFP peak bind to a promoter of PCG with accessible chromatin only in endoderm or mesoderm, both KZFP gene and PCG should be up-regulated. The potential target genes of KZFPs in differentiation from ESCs to endoderm or mesoderm were listed in supplementary table 8.

### The over- or under-representation analysis

This was performed using the method as previously described (D. Yang et al., 2012). Briefly, we first selected genes with BP term annotations. Subsequently, we filtered genes expressed during the differentiation of ESC into endoderm or mesoderm as the background. To analyse the over- or under-representation strengths of genes in each class relative to the background, we used the method based on hypergeometric distribution. The over- or under-representation strengths of each class were represented by -log (*p*) or log (*p*).

### Retrieving the metrics of functional constraint and evolutionary rate measurements

We characterized KZFP target genes using three different measurements of functional constraint: (1) the residual variation intolerance score (RVIS); (2) the probability of being intolerant to loss-of-function mutations (pLI); and (3) the selection against heterozygous loss of gene function (S_het_). We obtained the pLI and RVIS percentile from Ref.(Dickinson et al., 2016) and S_het_ scores from Ref.(Cassa et al., 2017). The number of synonymous substitutions per synonymous site (dS) and the number of nonsynonymous substitutions per nonsynonymous site (dN) between the ortholog pairs of human and chimpanzee were retrieved using BioMart from Ensembl database (Kinsella et al., 2011).

### Plasmids and siRNA

ZNF611 mammalian expression vector were constructed by PCR, followed by subcloning into pFLAG-CMV-2 (Sigma). The siRNA was purchased from Gene Pharma (Suzhou, China), and the following sequences were used. siZNF611-1: 5’-GCAGGUCAUCCCUUCAUUGTT-3’, siZNF611-2, 5’-GUGUAAUUCACUCCUGUCATT -3’. non-targeting siRNA, 5’-CCAUGUGAACUGGUCACCUTT -3’.

### Cell culture and transfections

The HEK293T cells were maintained in DMEM supplemented with 10% fetal bovine serum (Zhejiang Tianhang Biotechnology, Hangzhou, China).

The human ESC line H9 was cultured following the protocols as previously described (Ludwig et al., 2006). Briefly, H9 cells were plated as clumps in feeder-free conditions in six-well plates (Corning) coated with Matrigel (1: 80, BD Biosciences) in mTESR1 medium (Stem Cell Technologies). Gentle Cell Dissociation Reagent (STEMCELL Technologies) was used for passaging (1: 4-1: 7) every 4-7 days when cells reached 70-80% confluency on Matrigel. The differentiated cell clumps were manually removed before passaging to ensure high quality cultures.

When H9 cells reached 70 - 80 % confluency, the clumps were digested into single cells for optimal transfection efficacy. Cells were grown on 12-well plates coated with Matrigel. Single-cell passaging medium by adding Y-27632 (10 μM, Stem Cell Technologies) to mTeSR1 was used for the 1st day of feeder-free culture (Watanabe et al., 2007). Before the transfection, the differentiated cells were manually removed.

Both HEK293T cells and H9 human ESCs were transfected with Lipofectamine 2000 (Invitrogen, CA, USA) according to the manufacturer’s instructions.

### Quantitative PCR

Total RNA was isolated using Trizol kit (T9424, Sigma-Aldrich). Complementary DNA was synthesized using the cDNA synthesis kit (FSQ-101, TOYOTO, Osaka, Japan) according to the manufacturer’s instructions. Fluorescence real-time RT-PCR was performed using the KAPA SYBR® Faster Universal (KK4601, Biocompare, South San Francisco, CA, USA) and the ABI PRISM 7300 system (Perkin-Elmer, Torrance, CA). PCR was conducted in triplicates and standard deviations representing experimental errors were calculated. All data were analyzed using the ABI PRISM SDS 2.0 software (Perkin-Elmer). The expression of genes was normalized to the GAPDH gene. The PCR primers were listed in supplementary file 1.

### Chromatin immunoprecipitation (ChIP) assay

ChIP was performed as previously described (Yuan et al., 2015). The following primers were used: Primers: STK38 (5’-CAGCAAGCAACTCACCAGAG-3’, 5’-TCCTGTTGTCCTCACCCGTA-3’), BMPR2 (5’-TTTGTGTCTGGTCTGCTCGG-3’, 5’-GCACTTCCAGTGGCTCCG-3’). The percentage of immunoprecipitated DNA relative to the input was calculated and is shown as the mean ± SD from three independent experiments.

### Luciferase Assay

To generate the pGl3-STK38-P luciferase reporter,1.2 kb of DNA upstream the STK38 TTS promoter region with ZNF611 binding motif was amplified by polymerase chain reaction (PCR) and cloned into the Firefly luciferase reporter plasmid pGL3-Basic (Promega) using KPN1 and XHO1 restriction sites.0.7 kb of DNA upstream the STK38 TTS promoter region without ZNF611 binding motif was cloned into pGL3-Basic using same restriction sites to generate the pGl3-STK38-ΔP luciferase reporter. HEK293T cells seeded on 48-well plates were incubated until 60% to 70% confluence and then were transient transfected with 0.01 μg pGL3-STK38 plasmid using Lipofectamine 2000 (Invitrogen, CA, USA). After transfection 36-48 h, cell lysates were collected for Luciferase activity determination by Dual Luciferase Reporter assay systems (Promega). According to the manufacturer’s instructions, luciferase activity was normalized by Renilla activity to standardize transfection efficiency.

## Authors’ contributions

DY conceived, designed the study, revised the manuscript and supported the funding needed in this study. CT designed the experiments, partly supported the funding needed in this study and partly wrote the manuscript. FH partly designed the study, gave valuable suggestions, and partly supported the funding needed in this study. PS designed, carried out most of the analyses, and wrote the draft manuscript. QZ performed most of the experiments, and partly wrote the draft manuscript. AX, CG, YH participated in part of analyses. JL participated in part of experiments. All authors read and approved the final manuscript.

## Acknowledgements

Thanks for the support and help of bioinformatics platform of National Center for Protein Sciences (Beijing) in omics data processing.

## Competing interests

The authors declare that they have no competing interests.

## Notes

### Competing Interest Statement

The authors have declared no competing interest.

