## Supplementary file for "Novel non-transposable-element regulation patterns of KZFP family reveal new drivers of its rapid evolution"

(Supplementary Information)

#### Contents

**Supplementary Results**


- KRAB domains are evolutionarily young, but tend to be completely structured
- KZFPs, with older zinc fingers, contain variable and disordered KRAB domains
- Most of KZFPs tent to be highly structured
- Comparison of binding sites for KZFPs in this study and previous study
- KZFPs can play positive regulatory roles via binding to non-TE regions in ESCs and HEK293T cells


**Supplementary Methods**


- Data sources
- Primers used in quantitative PCR


**Supplementary References**

**Supplementary Tables (in the Excel files)**


Supplementary table 1. The structural disordered ratio and consecutive disordered region number of representative proteins (related to figure 2B&2C and supplementary figure 3-4).

Supplementary table 2. Supplementary table 2. The similarity values of the key amino acids in zinc fingers of each KZFP pairs (related to figure 2D).

Supplementary table 3. The percentage of peaks not overlapping with TEs (%) of each KZFP in multiple different genomic regions (related to figure 3).

Supplementary table 4. The comparison of binding bias of KZFPs with different data processing flows.

Supplementary table 5. The Spearman's rank correlation coefficients between divergence times of full protein, KRAB domain or zinc fingers and the expression level (related to figure 4).

Supplementary table 6. 80 species used in the sequence alignment of KRAB domains.

Supplementary table 7. The divergence time of the full protein, KRAB domain and zinc fingers in KZFPs.

Supplementary table 8. The potential target genes of KZFPs in the differentiation from ESCs to endoderm or mesoderm.

**Supplementary Figures**

Supplementary figure 1. The schematic diagram of the domain architecture of KZFP and the key amino acids in zinc finger binding to DNA.

Supplementary figure 2. The divergence time of full protein, KRAB domain and zinc finger in KZFPs.

Supplementary figure 3. The domain or protein disorder degree of KZFPs and other C2H2-ZFPs.

Supplementary figure 4. The disorder degree of KZFPs, other C2H2-ZFPs, TFs and other proteins.

Supplementary figure 5. The comparison of the accuracy between MACS2 and MACS1.4.

Supplementary figure 6. The binding bias of KZFPs across different zinc finger divergence time grades.

Supplementary figure 7. KZFPs can play positive regulatory roles via binding to non-TE regions in ESCs.

Supplementary figure 8. There are more KZFP peaks within open chromatin in ESCs than in HET293 cells.

Supplementary figure 9. The functions of KZFP target genes in the differentiation of ESC into endoderm.

Supplementary figure 10. The dynamic changes of expression levels of STK38 during the differentiation of hESC or mESC into mesoderm.

**Supplementary Results**

**KRAB domains are evolutionarily young, but tend to be completely structured**

Over half of human protein domains are ancient ones, existing across eukaryotes or even across cellular organisms. The protein domains originated after the origination of the common ancestor of chordata are regarded as evolutionarily young domains in this study. In our another study, we found that young domains tend to be highly disordered (Unpublished data). Disordered proteins and domains lack stable three-dimensional structures, but can perform important functions such as chaperones (van der Lee et al., 2014). Thus, KRAB domains, as definitely young domains, should be highly disordered according to the existing knowledge. To test this hypothesis, two parameters were used to measure the disorder degree of proteins and domains: structural disorder ratio (SDR) and the number of consecutive disordered regions (CDRN) (see Methods) in the protein or domain. Surprisingly, we found that most of KRAB domains (85.12 %, 309/363) are completely structured (SDR = 0) (supplementary fig. 3F), and SDR and CDRN of KRAB domains in KZFPs are obviously much lower than other domains in other C2H2 zinc finger proteins (C2H2-ZFPs) (supplementary fig. 3F&3G). The conformation of highly structured proteins and domains usually are rigid (Habchi, Tompa, Longhi, & Uversky, 2014), indicating that the functional mechanism of these proteins or domains tends to be single and changeless. Therefore, these results suggested that, in general, KRAB domain is a kind of rigid domain which is not easy to change its conformation, and its functional type is unique.

**KZFPs, with older zinc fingers, contain variable and disordered KRAB domains**

As a result, the KRAB domains in KZFPs containing older zinc fingers (Am-Th, *i. e.* divergence time grade from amniota to theria (over 159 Mya)) tend to have higher SDR values (supplementary fig. 3D). However, they do not contain significantly more CDRs (supplementary fig. 3E). In other words, these disordered residues are scattered in the KRAB domain and do not form a long continuous disordered region. This kind of disordered pattern can have strong adaptability and ensures that the overall conformation of the domain remains unchanged. For the 'Non-domain' region, mainly referring to the linker region between KRAB domain and C2H2 zinc fingers, the KZFPs containing old zinc fingers have both significant higher

SDR values (supplementary fig. 3D) and more CDRs (supplementary fig. 3E). This indicates that the linker region in old zinc-finger-containing KZFPs tend to be more flexible, making KZFPs have more variable conformation, which may be related to its diverse interaction pattern. Thus, these results suggested that part of KZFPs, with older zinc fingers, contain variable and disordered KRAB domains, responsible for diverse protein binding functions.

**Most of KZFPs tent to be highly structured**

Besides KRAB domain, the disorder degrees of C2H2 zinc fingers and non-domain regions in KZFPs are significantly lower than those in other C2H2-ZFPs (supplementary fig. 3F). Thus, it’s reasonable that the SDR of whole KZFP proteins are obviously lower than other C2H2-ZFPs. From the viewpoint of CDRN, KZFPs also contain less CDRs than other C2H2-ZFPs (supplementary fig. 3G). When analyzed within a larger scope, KZFPs also show much lower SDR and CDRN values compared with other transcription factors (TFs) or all of other proteins encoded by human genome (supplementary fig. 4A&4B; supplementary table 1). These results revealed that KZFPs are highly structured, suggesting that most of KZFPs are monotonous and unchangeable in structural conformation and functional mechanism.

**Comparison of binding bias for KZFPs in this study and previous study**

By analyzing binding sites of KZFPs, we found that over half of KZFPs tend to bind to non-TE regions in human genome (fig. 3A, supplementary table 3). However, two previous studies from the same lab obtained the opposite result (KZFPs tend to bind to TEs) based on the same ChIP-exo data(Helleboid et al., 2019; Imbeault, Helleboid, & Trono, 2017). To figure out the reason for this difference, we randomly selected the ChIP-exo data for 30 KZFPs (Imbeault et al., 2017). We considered the following possible differences: reference genome version (hg19 or hg38), mapping software (Bowtie2 or HISAT2), peak-calling software (MACS1.4 or MACS2) and the threshold of P value ($< 10^{-8}$ and $<10^{-16}$). Then we processed the data with different workflows based on different combinations of the above parameters, and found that peaks identified by MACS2 tend to bind to non-TE regions, while peaks identified by MACS1.4

tend to bind to TEs regardless of the changes of reference genome version, mapping software or P value (supplementary table 4), indicating the main reason for this difference is the different versions of MACS used in the ChIP-seq data processing.

We used MACS2, but MACS1.4 was used in the previous study (Helleboid et al., 2019; Imbeault et al., 2017). To determine which version of MACS is better, we compared the P values of peaks in 262 KZFPs identified by MACS1.4 and MACS2 on the basis that all other conditions are the same (we used hg38, HISAT2 and $10^{-8}$ as P value threshold). We observed regardless of peaks binding to TEs or non-TE regions, the P value of common peaks (identified by both methods) is smaller than the P value of unique peaks (identified by only MACS1.4 or MACS2) (supplementary fig. 5A). Then we counted the percentage of common peaks in peaks identified by MACS1.4 or MACS2 respectively, and found that about 85% peaks (averdivergence time in 262 KZFPs) in MACS2 are common peaks, whereas over 60% peaks (averdivergence time in 262 KZFPs) in MACS1.4 are unique peaks, indicating that the MACS2 is more accurate than MACS1.4 (supplementary fig. 5B). What's more, Tao Liu, the main developer of MACS program, also pointed that the precision and sensitivity of MACS2 are better than MACS1.4 (https://github.com/taoliu/MACS/issues/315). These results demonstrated that our result is more reliable.

**KZFPs can play positive regulatory roles via binding to non-TE regions in ESCs and HEK293T cells**

We examined the effects of overexpression of ZNF202, ZNF383, ZNF589, ZNF554, ZNF565 and ZFP14 on the expression of their predicted target genes in ESCs and HEK293T cells (supplementary fig. 7A-7C). When these KZFPs were overexpressed in ESCs respectively, all the predicted target genes were up-regulated (supplementary fig. 7D). However, when each of them was overexpressed in HEK293T cells, two target genes of ZNF202, PHC1 and ZBTB24, were up-regulated, and two target genes of ZNF565, TPK1 and UBE4A were also up-regulated, but the target genes of ZNF383 and ZNF554, including UBL3, CASD and SCAMP1, showed a significant down-regulated trend, and there was no change in the expression level of target genes of ZNF202, ZNF565 and ZFP14, including SEPSH1, UQCRH, AP2B1 and GRAMD1A (supplementary fig. 7E). To investigate whether the status of chromatin accessibility of target genes is associated with the positive regulation of KZFPs, we checked the chromatin accessibility of these target genes in ESCs and HEK293T cells. All KZFP peaks bound to non-TE regions in promoters of target genes within accessible chromatins in ESCs. However, KZFP peaks that bound to the promoters of up-regulated target genes were all within accessible chromatins in HEK293T cells, while KZFP peaks that bound to the promoters of down-regulated or unchanged target genes were all within inaccessible chromatins (supplementary fig. 7F). These observations indicated that KZFPs can play positive regulatory roles via binding to non-TE regions in ESCs and HEK293T cells, and suggested that the positive regulatory roles of KZFPs need their binding to accessible chromatin.

#### Supplementary Methods

##### Data sources

The data of RNA-seq, ChIP-seq, ChIP-exo, and ATAC-seq were downloaded from the public databases of GEO (Gene Expression Omnibus) (https://www.ncbi.nlm.nih.gov/geo/), ArrayExpress (https://www.ebi.ac.uk/arrayexpress/) and ENCODE (https://www.encodeproject.org/). The data set IDs were listed as following:

***RNA-seq data.*** Human early embryonic development (GSE72379, GSE101571); differentiation of hESCs into three germ layers (GSE17312); differentiation of hESCs into endoderm (E-MTAB-3158, GSE44875, GSE52657); differentiation of hESCs into mesoderm (GSE54968, GSE74665, GSE76523); differentiation of hESCs into ectoderm (GSE56152, GSE56796, GSE80264); human adult tissues/organs (E-MTAB-2836); developmental stage times from early organogenesis to adulthood in human (E-MTAB-6814).

***ChIP-seq or ChIP-exo data.*** ChIP-seq (GSE2523, GSE76494, ENCSR365GRX, ENCSR882ERE,

ENCSR449UFF, ENCSR157CAU, ENCSR000ECJ, ENCSR808FFI, ENCSR448UKK, ENCSR751PNN, ENCSR551KAP, ENCSR801BWR, ENCSR534NZI, ENCSR295GRB, ENCSR194BBY, ENCSR200JYP, ENCSR167KBO, ENCSR289NSN, ENCSR669GUM, ENCSR224NFP, ENCSR567XAM, ENCSR550ZZK, ENCSR595FAO, ENCSR897RVP, ENCSR167XFW, ENCSR538RDA, ENCSR776MDR, ENCSR011XCI, ENCSR603XLW, ENCSR017QBI, ENCSR878DES, ENCSR762DQP, ENCSR233MWH, ENCSR497IAS); ChIP-exo (GSE78099); differentiation of mESCs into mesoderm (GSE47948, GSE67868).

***ATAC-seq data.*** Differentiation of hESCs into endoderm (GSE97992); differentiation of hESCs into mesoderm (GSE85881); HEK293T cells (GSE121252).

**Primers used in quantitative PCR**

qPCR primers

| Gene name | Forward | Reverse |
|---|---|---|
| AP2B1 | 5’-CTCTTTCCAGACGTAGTGAACTG-3’ | 5’-GGAGCGGCTCACAGAGATATT-3’ |
| BMPR2 | 5’-CGGCTGCTTCGCAGAATCA-3’ | 5’-TCTTGGGGATCTCCAATGTGAG-3’ |
| CSAD | 5’-CTTCTCCAGGATACCTCGAACC-3’ | 5’-CAGAGCCACATCGTAGAACTTG-3’ |
| GAPDH | 5’-GGAGCGAGATCCCTCCAAAAT-3’ | 5’-GGCTGTTGTCATACTTCTCATGG-3’ |
| GRAMD1A | 5’-ATAAGCAGCGTAATGAGGACTTC-3’ | 5’-AGGGCGCAGGAGTAATCCA-3’ |
| PHC1 | 5'-TTCCACCAATGGGAGTTCTAGC-3' | 5'-GCACTGCTTGTCGTTCATAAAGT-3' |
| SCAMP1 | 5’-TTTCGACAGTAACCCGTTTGC-3’ | 5’-TTGGGTACATTAGGCATCTTCAC-3’ |
| SEPHS1 | 5'-ACTTGTGTCATTCCTTTGAGGC-3' | 5'-CCCATCATGTAAGGGTCGTCTA-3' |
| STK38 | 5’-ACAAAGGAAAGGGTGACAATGAC-3’ | 5’-CTTCCGAGCATGTGCTGATCT-3’ |
| TPK1 | 5'-AGCCTTTGGACAACTATTTTCGT-3' | 5'-AGCTCACATCCCTTAGTAGCA-3' |
| UBE4A | 5'-AACATCTCAAGTAACCCCTTTGC-3' | 5'-CACGCTATTATCCGAGTCATCTG-3' |

| | | |
|---|---|---|
| UBL3 | 5'-AGTAATGTCCCGGCGGATATG-3' | 5'-TCAGAAGCAGAATCGTTAGGAGA-3' |
| UQCRH | 5'-GAGGACGAGCAAAAGATGCTT-3' | 5'-CGAGAGGAATCACGCTCATCA-3' |
| ZBTB24 | 5'-AGAAACATCTGCGAACACAC-3' | 5'-GTTGTCTAAGCGAGCAAAC-3' |
| ZFP14 | 5'-GCAACTTCATTTCACTAGGACCT-3' | 5'-GGCAGTATCTTCTTGTCCCTTC-3' |
| ZNF202 | 5'-GACCAGAGAGGCGGACAAAG-3' | 5'-GCCGTGAACATGGACAGTCA-3' |
| ZNF383 | 5'-ATGGCTGAGGGATCAGTGATG-3' | 5'-GAAACCAGATTGCCGTAGTTCT-3' |
| ZNF554 | 5'-CTGAGCCCAGCCCTTGTTTTA-3' | 5'-TGGTGGTTTAAGACTGAATCGG-3' |
| ZNF565 | 5'-TTGGCCTCACTAGGACTCTCC-3' | 5'-CCCAGTTGAATGCCCCGATTT-3' |
| ZNF589 | 5'-GCAGAAGATCAACGAGTGGAA-3' | 5'-CCAAACGTGACTGCTCCAAAC-3' |
| ZNF611 | 5’-TAGGCCCCAAACCCAGATTTC-3’ | 5’-TGCAAGGTATTGCTCGTGATT-3’ |

**Supplementary figure legends**

**Supplementary figure 1. The schematic diagram of the domain architecture of KZFP and the key amino acids in zinc finger binding to DNA.** All KZFPs contain a KRAB domain and a C-terminal C2H2 zinc finger array with DNA-binding potential (at positions 6, 3 and -1). Some KZFPs contain other domains, such as SCAN domain and DUF3669 domain.

**Supplementary figure 2. The divergence time of full protein, KRAB domain and zinc finger in KZFPs. A,** the cumulative curve of divergence time of full protein, KRAB domain and zinc finger in KZFPs. **B**, the venn diagram of the number of KZFPs with the divergence time of Eutheria**.** **C**, the over- or under-representation strength of KZFPs containing variant KRAB domains (vKRAB) at three different KRAB domain divergence time grades. Numbers in the heatmap represent the percentage of KZFPs containing vKRAB or KZFPs containing standard KRAB domains (sKRAB) in three different KRAB domain divergence time grades. The over- or under-representation strengths of each class were represented by -log (p) or log (p), respectively and were shown in the heat map.

**Supplementary figure 3. The domain or protein disorder degree of KZFPs and other C2H2-ZFPs.** A, the schematic diagram of structure of KZFPs and other C2H2-ZFPs. **B&C**, the domain or protein SDR **(B)** or CDRN **(C)** between KZFPs containing variant KRAB domain (vKRAB) and KZFPs containg standard KRAB domain (sKRAB). **D-E**, the domain or protein SDR **(D)** or CDRN **(E)** in KZFPs with different zinc finger age grade. **F-G**, the SDR **(F)** or CDRN **(G)** of the domains within KZFPs and other C2H2-ZFPs and SDR values of the whole proteins. The values of upper and lower quartiles, and the median are indicated the same as **figure 1**. The differences of CDRN between different categories are examined by Mann–Whitney U test. The corrected P values are shown in the top of each panel. The abbreviations in the figure: Am-Th: Amniota-Theria, Eu-Ha: Eutheria-Haplorrhini, Si-Ho: Simiiformes-Homo.

**Supplementary figure 4. The protein disorder degree and CDRN of KZFPs, other C2H2-ZFPs, TFs and other proteins.** A, the SDR values of KZFPs, other C2H2-ZFPs, TFs and other proteins. **B**, the CDRN of KZFPs, other C2H2-ZFPs, TFs and other proteins. The values of upper and lower quartiles, and the median are indicated the same as **figure 1**. The differences of CDRN between different categories are examined by Mann–Whitney U test. The corrected P values are shown in the top of each panel.

**Supplementary figure 5. The comparison of the accuracy between MACS2 and MACS1.4. A**, box-plots of -lg (*p*) of common peaks called by both MACS2 and MACS1.4 and unique peaks called only by MACS2 or MACS1.4. **B**, the percentage of common peaks and unique peaks in all peaks called by MACS2 or MACS1.4. Peak were divided into four types: Common-TE (common peaks binding to TEs), Common-nonTE (common peaks binding to non-TE regions), Unique-TE (unique peaks binding to TEs) and Unique-nonTE (unique peaks binding to non-TE regions). The genome version hg38, HISAT2 (mapping tools), and P value threshold ($10^{-8}$) were used in this analysis. The differences of -lg (*p*) between different categories are examined by Mann–Whitney U test. The corrected P values are shown in the top of each panel. **: $p < 0.01$.

**Supplementary Figure 6. The binding bias of KZFPs across different zinc finger divergence time grades. T**he binding bias of KZFPs with different zinc finger divergence time grades in genome, intergenic region, all gene, PCG, non-PCG, all intron, intron in PCG or non-PCG, all exon, exon in PCG or non-PCG, 5'UTR, 3' UTR and PCG promoter. The values of upper and lower quartiles, and the median are indicated the same as **figure 1**. The number of KZFPs in each box can be found in Supplementary table6. The differences of the percentage of peaks not overlapping with TEs (%) between different categories are examined by Mann–Whitney U test. The corrected P values are shown in the top of each panel. The abbreviations in the figure: Am-Th: Amniota-Theria, Eu-Ha: Eutheria-Haplorrhini, Si-Ho: Simiiformes-Homo.

**Supplementary figure 7. KZFPs may play positive regulatory roles via binding to non-TE regions in ESCs. A&B**, the changes of KZFPs at mRNA level upon overexpression of corresponding KZFPs (ZNF202, ZNF383, ZNF589, ZNF554, ZNF565 and ZFP14) in ESCs (**A**) and HEK293T cells (**B**). **C**, the changes of KZFPs at protein level upon overexpression of corresponding KZFPs by western blot in HEK293T cells. Western blot using an anti-Myc antibody. **D&E**, the changes of target genes at mRNA level upon overexpression of corresponding KZFPs in ESCs (**D**) and HEK293T cells (**E**). Total RNAs from KZFPs transfected HEK293T or ES cells were subjected to real-time quantitative PCR (qPCR) analysis. Relative mRNA levels of predicted target genes were normalized to GAPDH. The ratio values (relative expression level / the averdivergence time control value) were shown. The data are represented as the mean ± S.D. (n = 3). Control: empty vector. **F**, binding regions of KZFPs (ZNF202, ZNF383, ZNF589, ZNF554,

ZNF565 and ZFP14) and the chromatin accessibility at genomic loci corresponding to target genes in HEK293T cells, ESCs and endoderm or mesoderm. The differences of expression level between control and KZFP overexpression are assessed by t-test. **: $p < 0.01$, *: $p < 0.05$.

**Supplementary figure 8. More KZFP peaks within open chromatin in h1 ESCs than in HET293 cells.** **A&B**, box-plots for percentage of KZFP peaks binding to TEs or non-TE regions within different regions in genome (**A**) or PCG promoter (**B**). According to chromatin accessibility in h1 ESCs and HEK293T cells, the different regions are composed of four parts: accessible only in ESCs, accessible only in HEK293T cells, accessible in both cell types and inaccessible in both cell types. The values of upper and lower quartiles, and the median are indicated the same as **figure 1**. The differences of percentage between different categories are examined by Mann–Whitney U test. The corrected P values are shown in the top of each panel.

**Supplementary figure 9. The functions of KZFP target genes in the differentiation of ESC into endoderm.** **A**, the significantly over- or under-represented biological process terms for the two types of PCGs (KZFP-TE PCGs and KZFP-nonTE PCGs) in the differentiation of ESC into endoderm. The over- or under-representation strengths of each class were represented by -log (p) or log (p), respectively and were shown in the heat map. **B**, the tolerance to functional variants between KZFP-TEs and KZFP-nonTEs in differentiation of ESC into endoderm. **C**, the evolutionary rate of KZFP-TEs and KZFP-nonTEs in differentiation of ESC into endoderm. For the box plots (**B&C**), the values of upper and lower quartiles, and the median are indicated the same as **figure 1**. The differences of the tolerance to functional variation and the evolutionary rate between different categories are examined by Mann–Whitney U test. The corrected P values are shown in the top of each panel.

**Supplementary figure 10. The dynamic changes of the expression levels of STK38 during the differentiation of hESC or mESC into mesoderm.** **A**, the dynamic changes of STK38 expression level during differentiation of hESC into mesoderm in 3 datasets. **B**, the dynamic changes of STK38 expression level during differentiation of mESC into mesoderm in 2 datasets. TPM, transcripts per million reads.

**Supplementary fig. 1**

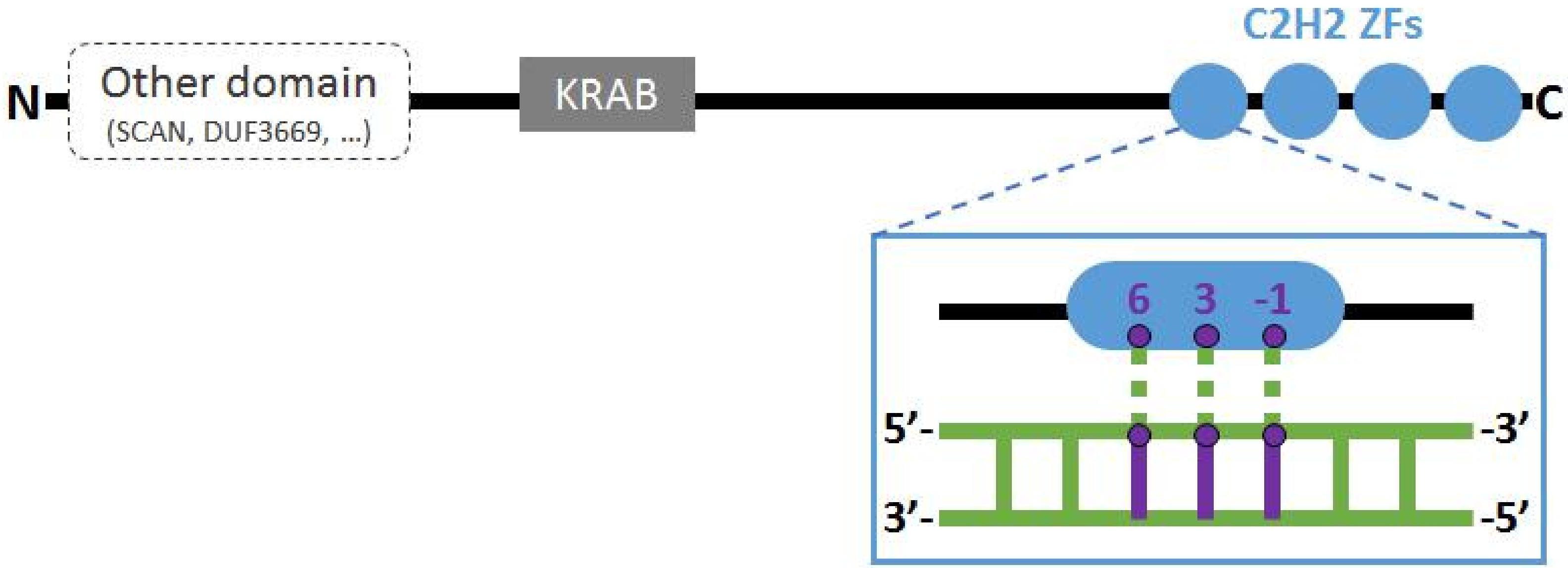

**Supplementary fig. 2**

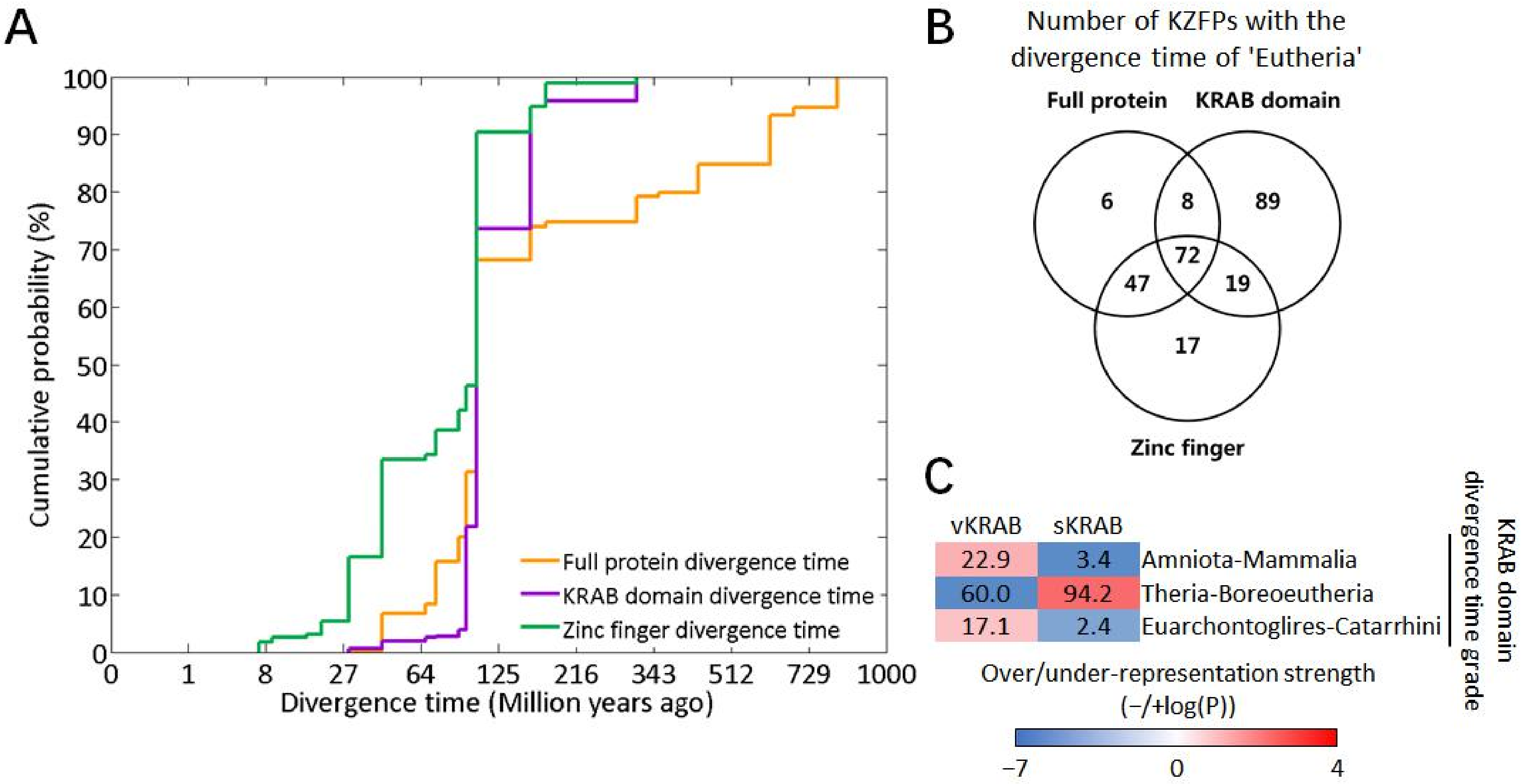

### Supplementary fig. 3

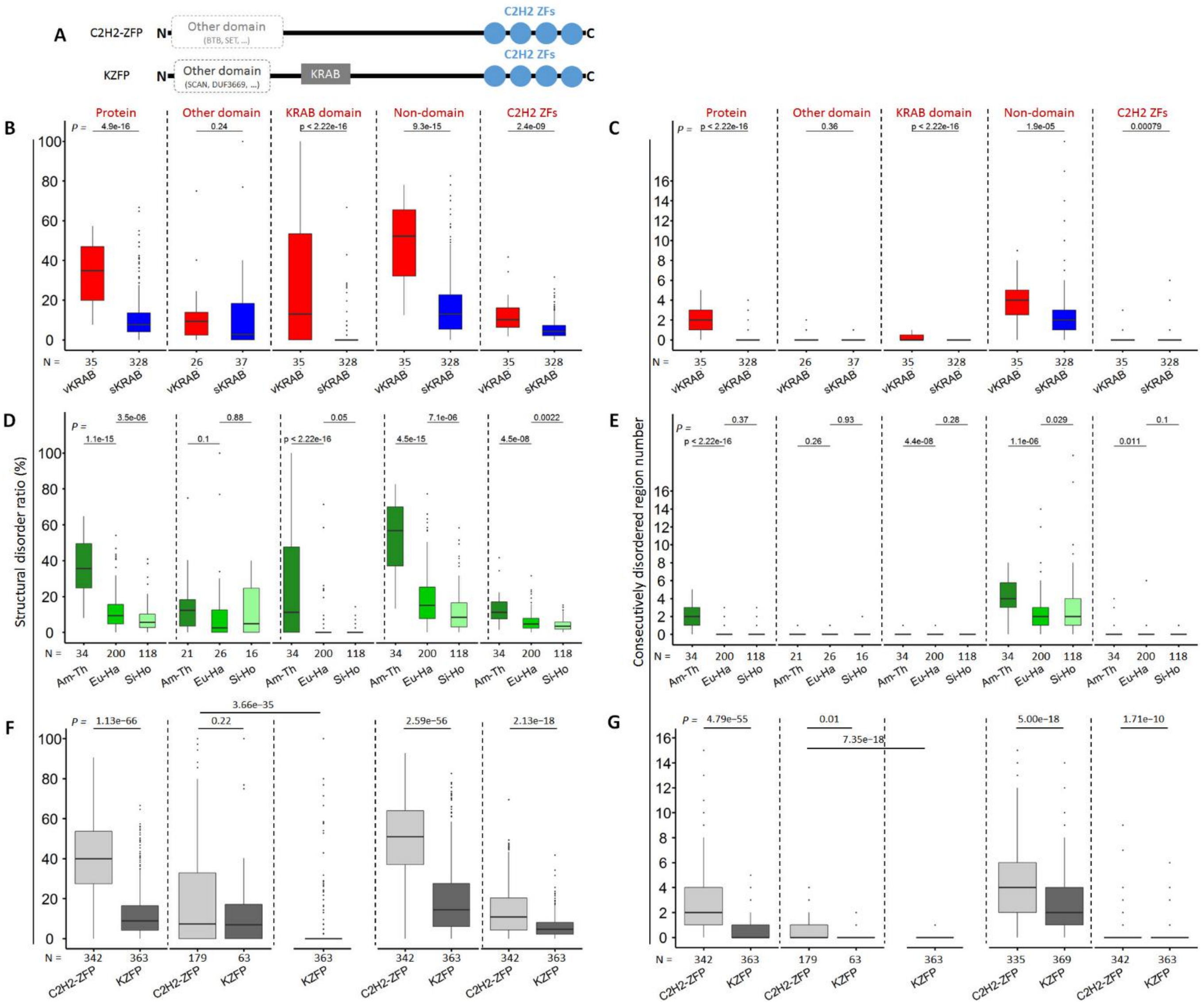

**Supplementary fig. 4**

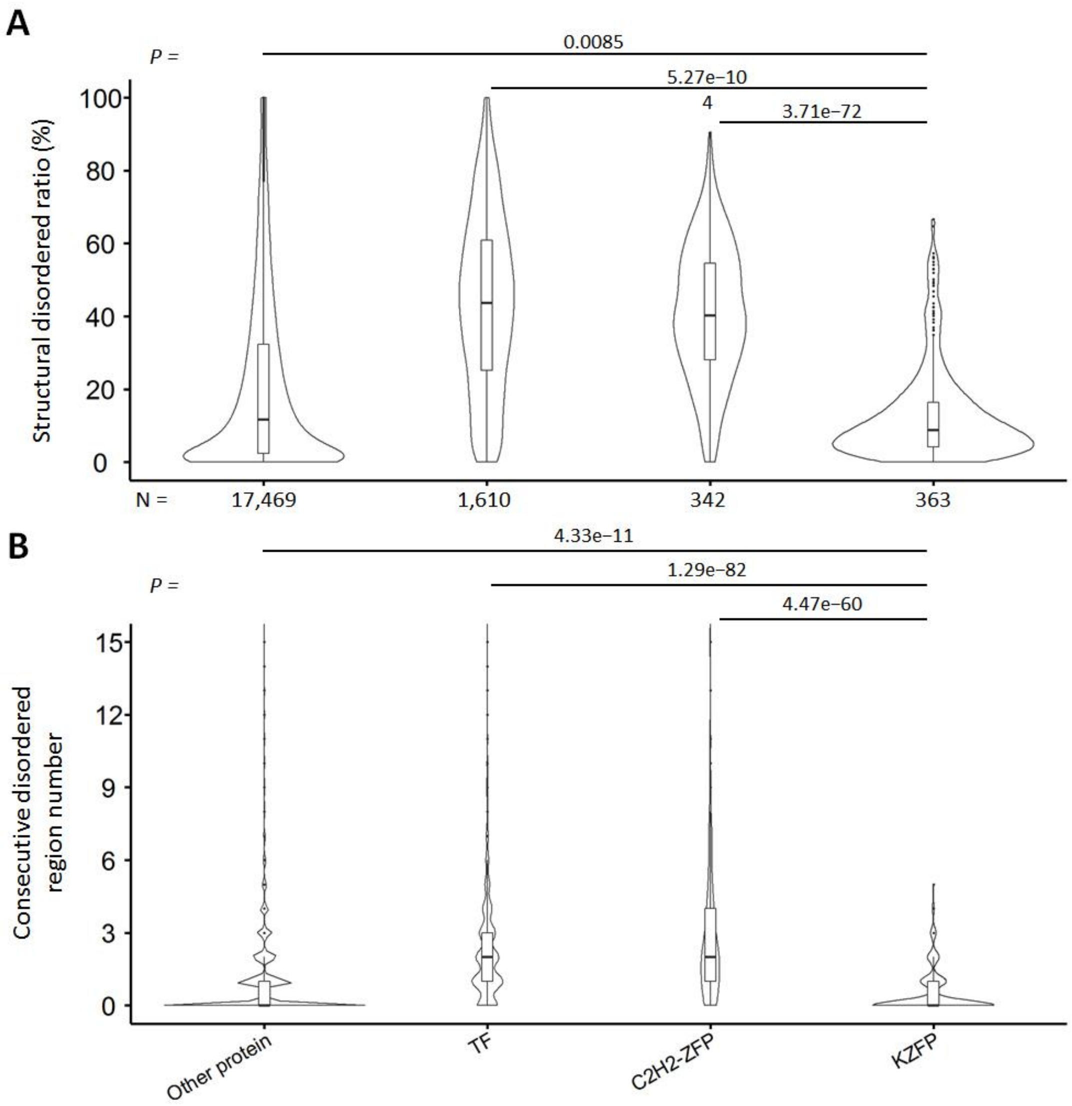

**Supplementary fig. 5**

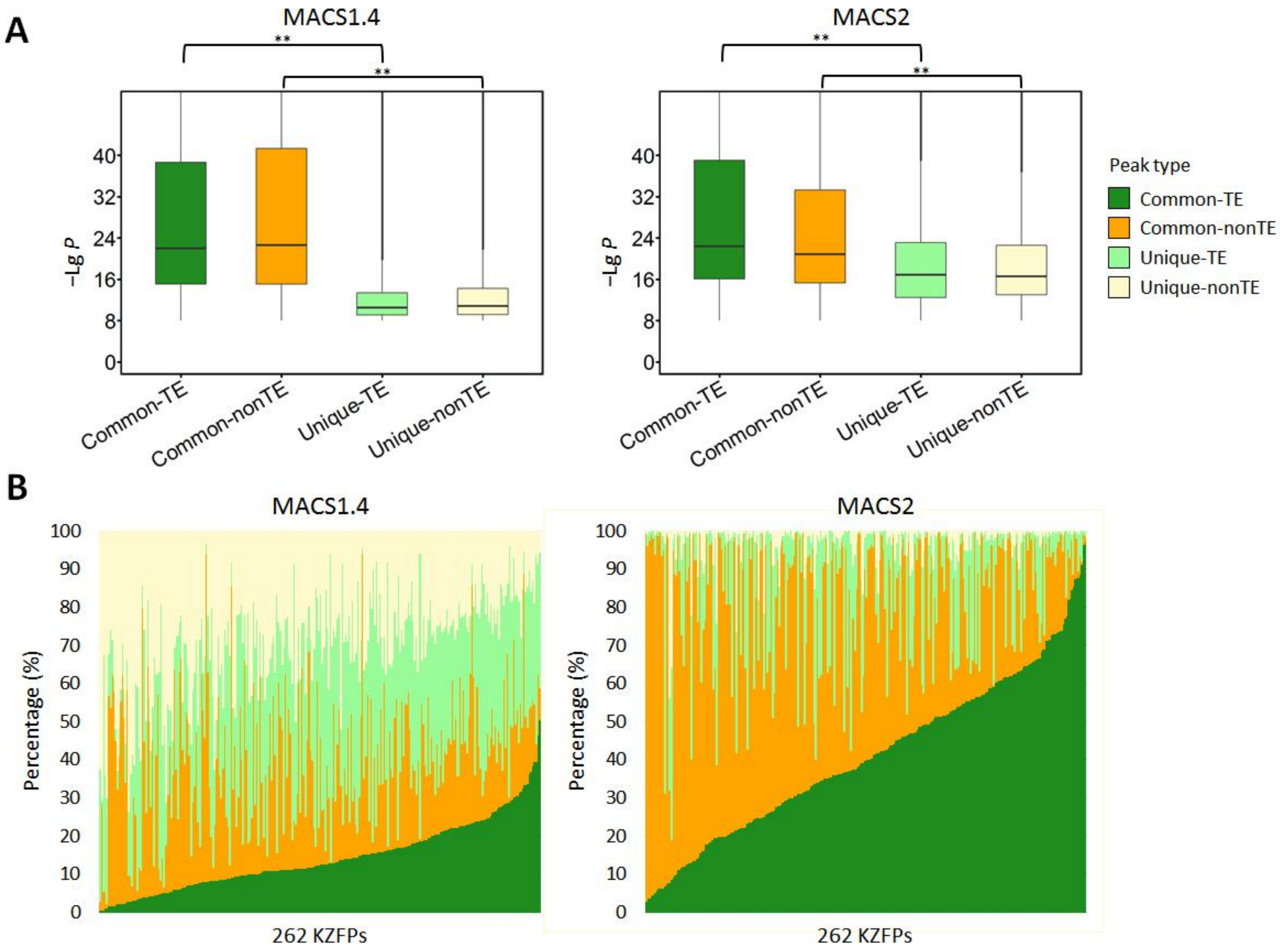

**Supplementary fig. 6**

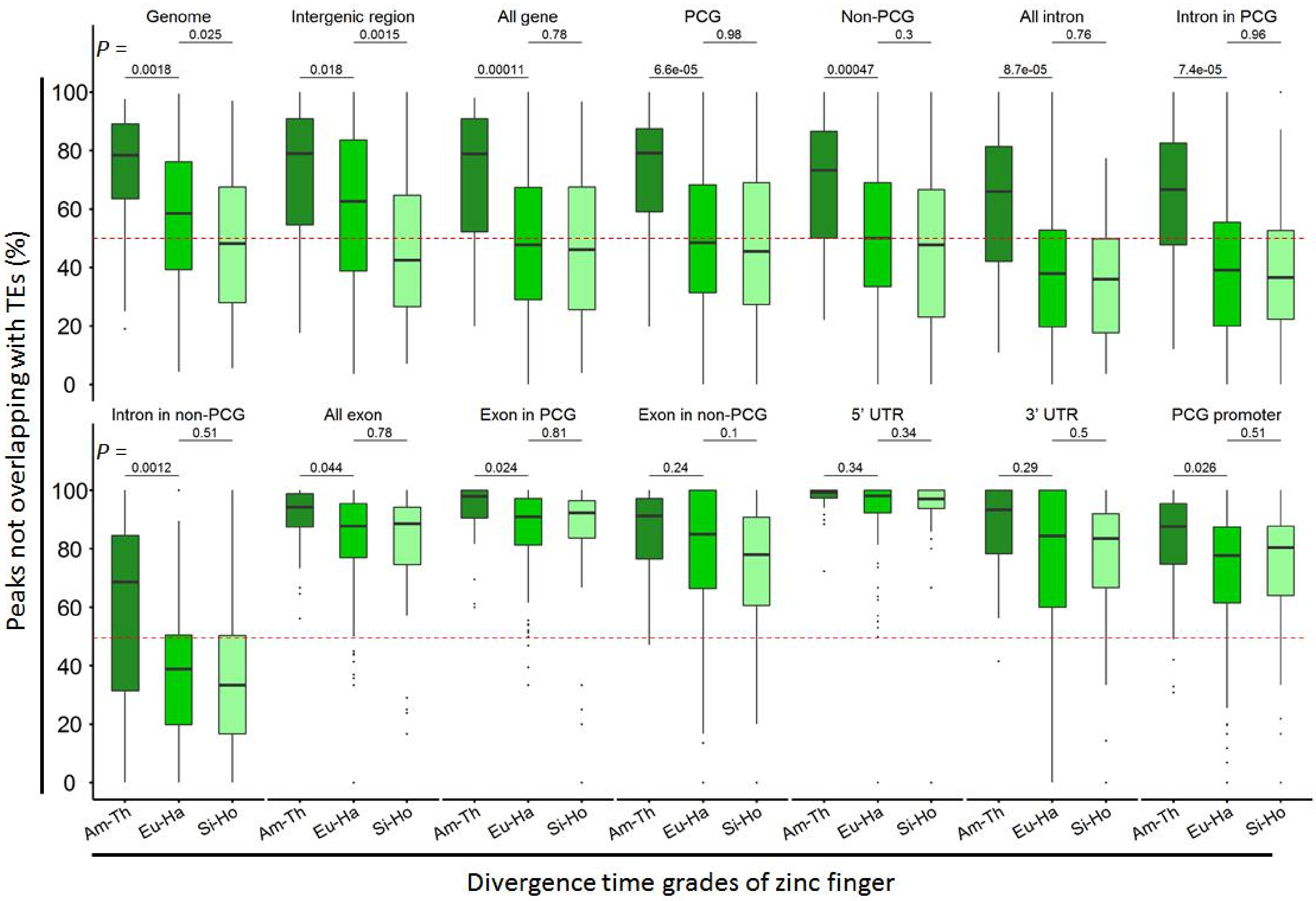

#### Supplementary fig. 7

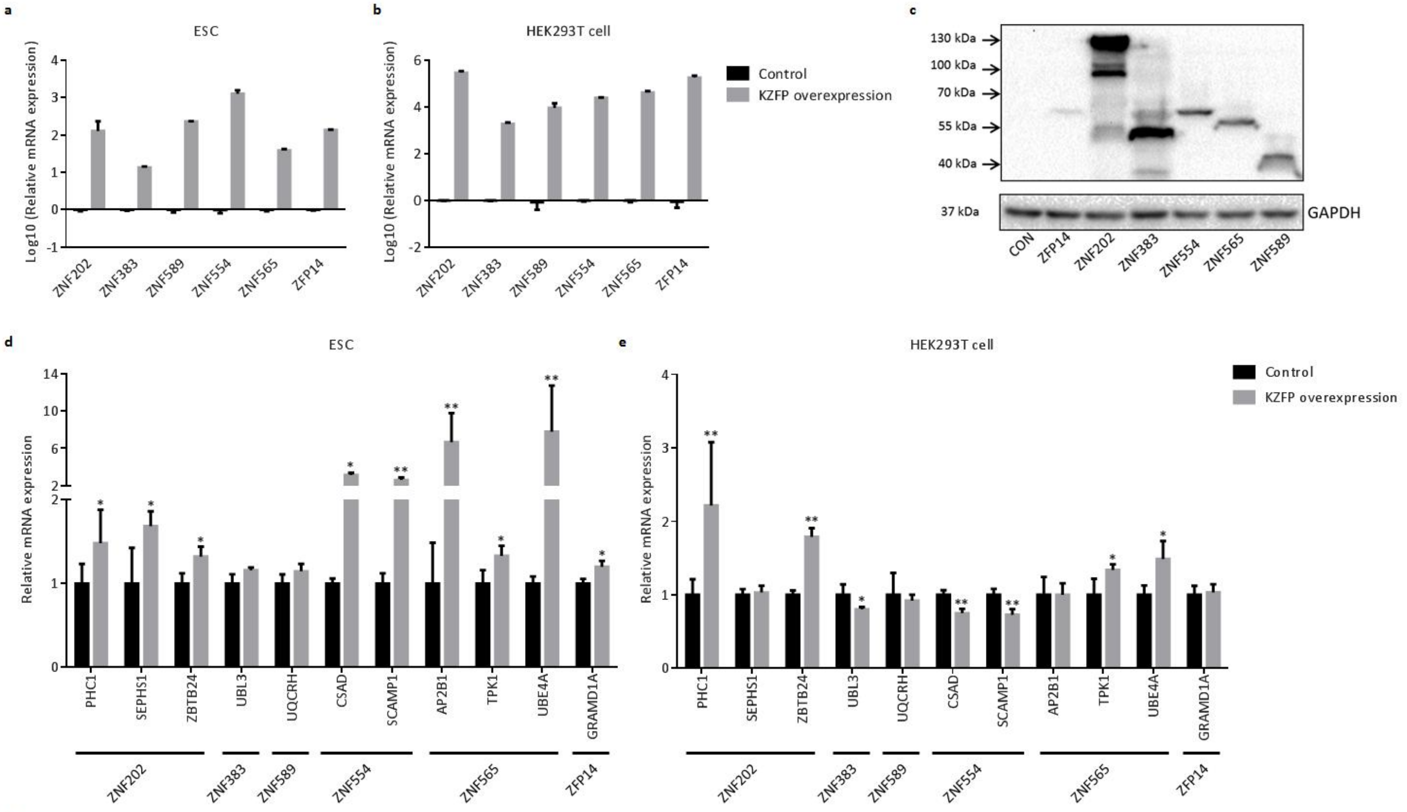


f

| Differentiation process | KZFP name | Target gene name | KZFP binding region | Chromatin accessibility: ESC | Chromatin accessibility: Endoderm or mesoderm | Chromatin accessibility: HEK293T cell | Experimental results of regulation trend: ESC | Experimental results of regulation trend: HEK293T cell |
|---|---|---|---|---|---|---|---|---|
| ESC→ Endoderm | ZNF202 | PHC1 | Non-TE | accessible | inaccessible | accessible | Positive | Positive |
| | | SEPHS1 | Non-TE | accessible | inaccessible | inaccessible | Positive | No change |
| | | ZBTB24 | Non-TE | accessible | inaccessible | accessible | Positive | Positive |
| | ZNF383 | UBL3 | Non-TE | accessible | accessible | inaccessible | Positive trend | Negative |
| | ZNF589 | UQCRH | Non-TE | accessible | inaccessible | inaccessible | Positive trend | No change |
| ESC→ Mesoderm | ZNF554 | CASD | Non-TE | accessible | accessible | inaccessible | Positive | Negative |
| | | SCAMP1 | Non-TE | accessible | accessible | inaccessible | Positive | Negative |
| | ZNF565 | AP2B1 | Non-TE | accessible | accessible | inaccessible | Positive | No change |
| | | TPK1 | Non-TE | accessible | accessible | accessible | Positive | Positive |
| | | UBE4A | Non-TE | accessible | accessible | accessible | Positive | Positive |
| | ZFP14 | GRAMD1A | Non-TE | accessible | accessible | inaccessible | Positive | No change |

**Supplementary fig. 8**

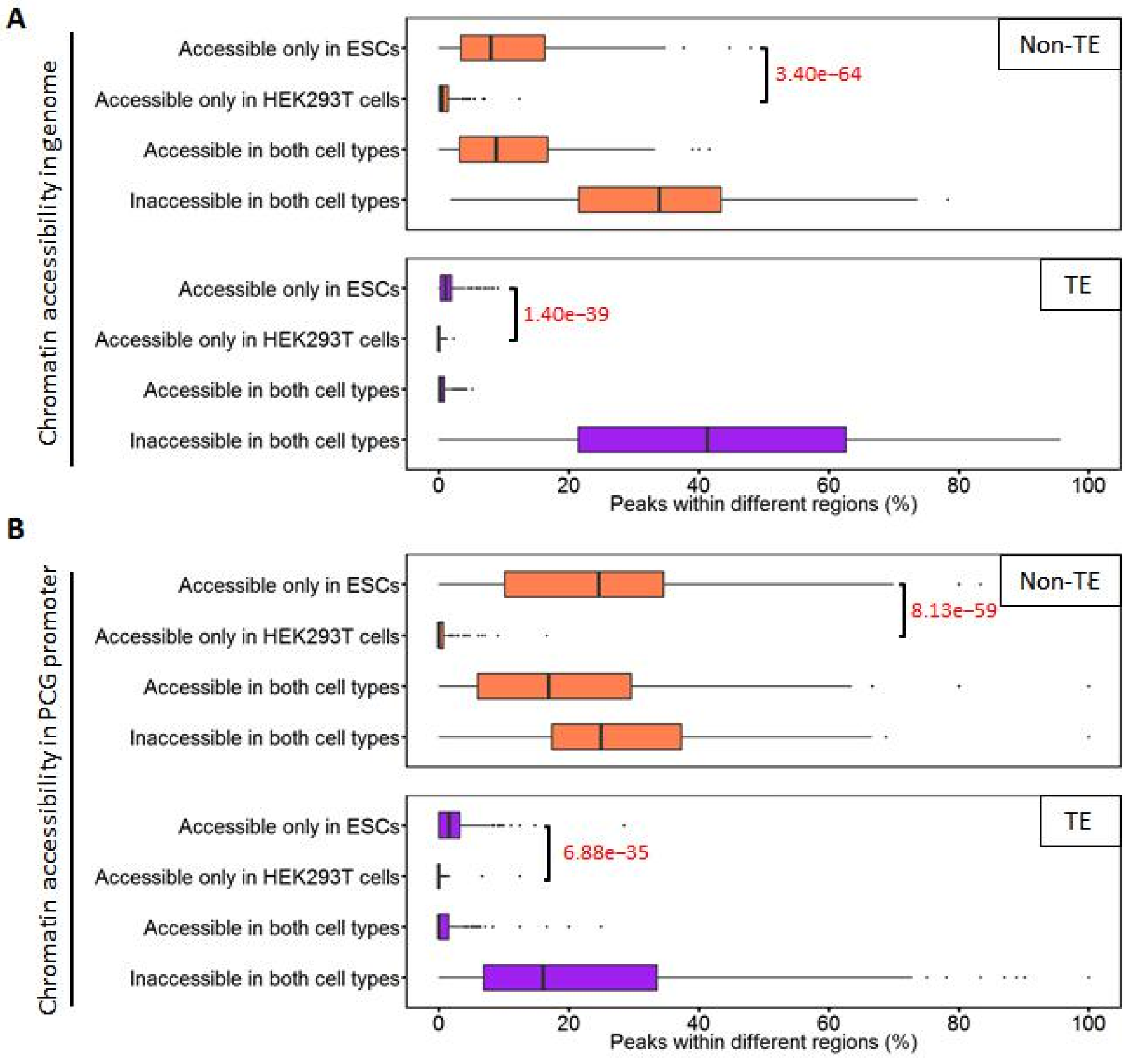

**Supplementary fig. 9**

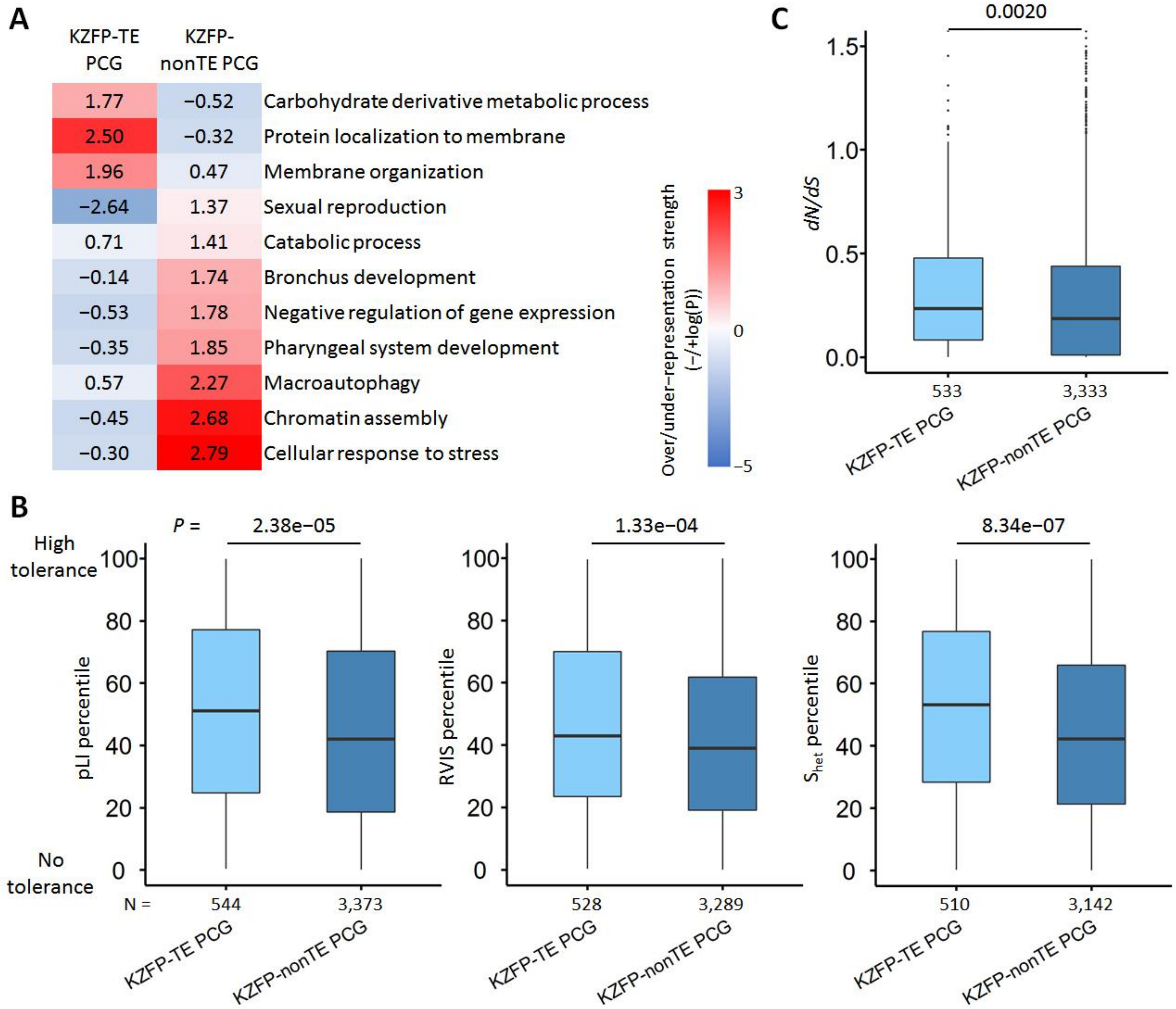

**Supplementary fig. 10**

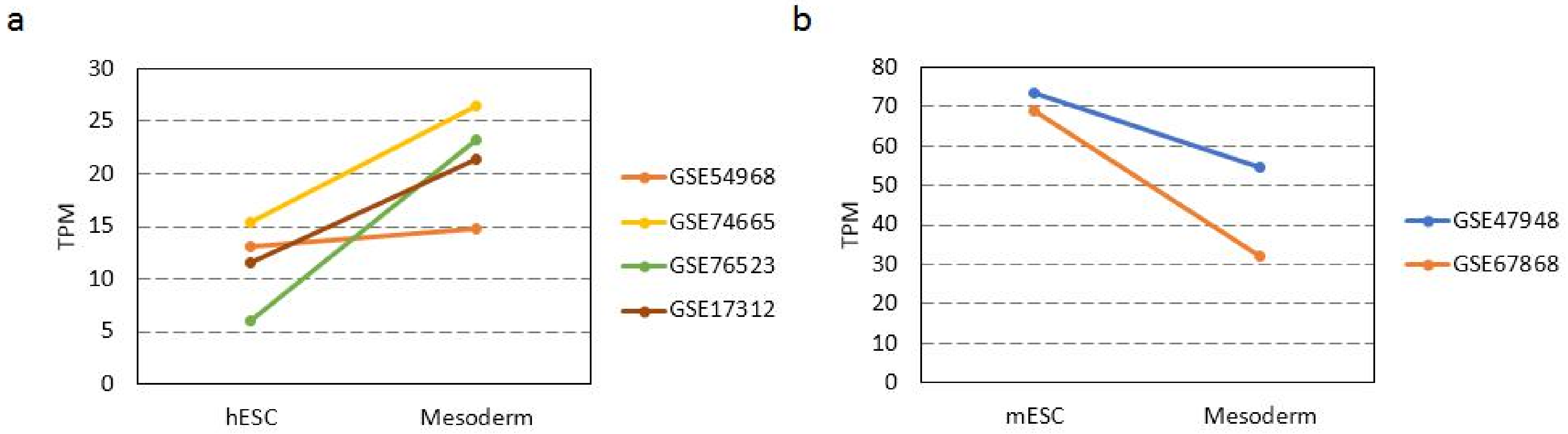
